# Migration strategy and site fidelity of the globally threatened Sociable Lapwing *Vanellus gregarius*

**DOI:** 10.1101/2020.03.31.017848

**Authors:** Paul F. Donald, Johannes Kamp, Rhys E. Green, Ruslan Urazaliyev, Maxim Koshkin, Robert D. Sheldon

## Abstract

Population declines of the critically endangered Sociable Lapwing are probably due to high mortality along its migration routes or on its wintering grounds, both of which are very poorly known. We therefore undertook a long-term study of the species’ movements using satellite tagging, colour-ringing and targeted field surveys. We also compiled a database of historical and recent sightings of the species from published and unpublished sources. There were two migration flyways from the breeding grounds in Kazakhstan, along which birds used different staging strategies. A longer western route (c. 5200 km) takes birds west to southern Russia, then south through the Caucasus and the Levant to wintering areas in Saudi Arabia and eastern Sudan. A shorter eastern route (c. 2800 km) takes birds south to Turkmenistan and Uzbekistan, then over the mountains of northern Afghanistan to wintering areas in Pakistan and north-western India. In spring, birds of the western flyway cut out the Caucasus by making a direct crossing of the Caspian Sea from Azerbaijan. The migration strategy is characterised by infrequent long-distance movements followed by often lengthy stopovers in a small number of staging areas that are used consistently across years, and by high individual and low between-individual consistency in patterns of movement, both spatially and temporally. At least four main autumn stopover areas and one additional spring stopover area were identified along the longer western route, but only one autumn and one spring staging area along the eastern route. There was no relationship between latitude or longitude of capture for tagging or colour ringing and the subsequent migration route used, and the same breeding colonies could contain breeding adults and produce chicks of both flyway populations, suggesting that no clear migratory divide exists within the breeding range. Sociable Lapwings spend around a third of the year on their breeding grounds, a third on their wintering grounds and a third moving between them. Birds were highly faithful to their passage and wintering sites, but showed low fidelity to breeding sites. The migration stopover areas and the wintering sites are usually located at the interface of agriculture, particularly irrigated cropland along rivers, and dry steppe-like or desert habitats. This species selects, and perhaps relies upon, agricultural habitats throughout its entire life cycle, and its heavy reliance on some of the world’s most anciently cultivated regions suggests that this synanthropic relationship may have evolved over many millennia. The recent emergence of irrigated cropfields in Arabia is likely to have allowed birds using the western route to winter well north of their previous wintering range and maybe to spread into new wintering areas along the coasts of the Arabian Gulf. The concentration of large numbers of birds at a small number of traditional but unprotected migration stopover areas offers the opportunity to quantify and monitor the global population size, for which we derive a tentative estimate of c. 24,000 individuals (95% CL: 13,700 – 55,560). However, it also makes the species particularly vulnerable to hunting and small-scale habitat change. Illegal hunting along the western flyway is identified as the most plausible driver of the species’ decline.

## INTRODUCTION

Sociable Lapwing *Vanellus gregarius* is a migratory shorebird that is listed on the IUCN Red List as Critically Endangered on the basis of severe population declines across its range (Eichhorn & Khrokov 2002, Sheldon *et al.* 2012, BirdLife International 2020). The species once bred from Ukraine (Kasparek 1992) to western China (Kamp *et al.* 2010), and small remnant populations may survive in southern Russia, but the breeding range is now almost entirely restricted to the steppes of central and northern Kazakhstan where post-Soviet changes in agriculture are profoundly altering the landscape (Kamp *et al.* 2011, Sheldon *et al.* 2012). Here, post-Soviet changes in steppe management and declines in the populations of natural grazers, particularly Saiga *Saiga tatarica*, have led to the species becoming largely confined to the periphery of villages that maintain sizeable herds of livestock, which in the absence of native grazers keep the steppe grazed to a level suitable for the species to nest (Kamp *et al.* 2009). Research on the species’ demography suggested that current levels of productivity are not particularly low, and that low adult survival is more likely to be the demographic driver of recent population declines (Sheldon *et al.* 2013). This, together with illegal hunting of birds migrating south in autumn through the Caucasus and Middle East (e.g. Murdoch & Serra 2006, Hofland & Keijl 2008, Brochet *et al.* 2016, Brochet *et al.* 2019a, Brochet *et al.* 2019b), suggests that the factors driving the species’ decline are likely to be acting as birds migrate from their breeding grounds to their wintering areas, or on the wintering grounds themselves.

Prior to this study, what little was known about the species’ wintering areas came from opportunistic sight records and the collection localities of museum specimens (Cramp 1983, Urban *et al.* 1986, BirdLife International 2001), which indicated that they lay in east Africa, particularly eastern Sudan, and northern Ethiopia and Eritrea, and in Pakistan and north-western India. However, the state of knowledge was poor; prior to this study the most recent sight record of the species in Sudan was before 1950, and the small number of records reported from India each year were insufficient to account for more than a tiny proportion of even the most pessimistically estimated population. Furthermore, the routes taken by birds to reach these areas were almost entirely unknown. This greatly hampered efforts to assess threats to the species outside its breeding season. We therefore undertook a long-term study of the species’ migration patterns and wintering areas using satellite tagging, colour ringing and targeted field surveys. We also collected sightings and specimen records, both historical and recent, and georeferenced these. Satellite tracking led to the identification of a number of areas that appeared to be used consistently by tagged birds on migration, and subsequent field visits to some of these sites confirmed that birds were present in sometimes large numbers. In particular, large number of birds have been seen in autumn along the Manych river in the Stavropol region of south-western Russia (Field *et al.* 2007), on irrigated agricultural land in southern Turkey along the border with Syria (Biricik 2009), and in spring in northern Syria in fields and steppe-like grazed grassland (Murdoch & Serra 2006, Hofland & Keijl 2008, Asswad 2014). In 2015 the world’s largest aggregation in recent years was discovered in 2015 at a site known as Tallymarzhan (also Tallymerjen), which straddles the border between eastern Turkmenistan and south-western Uzbekistan (Donald *et al.* 2016, Azimov *et al.* 2018). Here we present the first systematic overview of this species’ migration strategy, placing previous publications on the numbers of birds at individual sites in a flyway context and identifying a number of as yet un-surveyed areas where important aggregations of birds may occur. We also assess for the first time patterns of site fidelity on both the breeding and wintering grounds, and identify broad patterns of habitat selection.

## METHODS

### Satellite tagging

Between 2007 and 2015 we trapped 29 Sociable Lapwings on their breeding grounds in central and eastern Kazakhstan (three in 2007, two in 2008, eight in 2010, two in 2011, three in 2013, three in 2014 and eight in 2015). All but five birds were tagged within 70 km of the village of Korgalzhyn in central Kazakhstan, the remaining five were tagged in eastern Kazakhstan between the towns of Semey (Semipalatinsk) and Qalbatau, between 750 and 870 km east of Korgalzhyn. Birds were caught initially using walk-in cage traps placed over nests, thereby catching mostly females (since males rarely incubate), but in 2015 we switched to using leg nooses and mist nets set over a model owl near adults with chicks, in order to try to minimise disturbance at the nest and to catch more males. Nevertheless, our sample remains heavily weighted towards females (21 of 29 tagged birds). The first five birds were fitted with 9-g Argos PTTs, thereafter all birds were fitted with 5-g Argos PTTs (both supplied by Microwave Telemetry Inc.). All tags were set to a duty cycle of 10 hours on followed by 48 hours off. Locations were estimated using the Doppler effect and each was assigned an accuracy code (in order of decreasing accuracy: 3, 2, 1, 0, A, B and Z). The tags were fitted to birds using a modified Rappole-Tipton leg-loop harness (Rappole & Tipton 1991) made from silastic tubing (Dow-Corning Inc.). Tagged birds were also fitted with unique combinations of colour rings.

### Processing satellite tracking data

The distance between successive locations of tagged birds was estimated using the haversine formula, which estimates great-circle distances, and the direction of travel estimated as the forward azimuth of movement along a great-circle route. Due to (1) largely un-quantifiable inaccuracies in the Argos locations (Boyd & Brightsmith 2013), (2) the high proportion of locations in our data that fell in low classes of accuracy, and (3) the often long intervals between successive locations, particularly in the early years of the study and when birds were on the move, the satellite-tracking data were incapable of estimating small-scale movements with any degree of confidence. We therefore use them only to make inferences about large scale (>100 km) movements and the approximate timing of migration events. We attempted to distinguish genuine movements from random measurement error by applying a number of filters, in addition to the Kalman filtering that was automatically applied to all data after 2011 (Lowther *et al.* 2015). We plotted all tracks in GIS and connected fixes with lines using the ET GeoWizards extension in ArcGIS Pro to identify and remove all obvious major outliers, checking that the correct Argos location solution had been selected in each case. For all apparent movements of over 20 km we then compared the haversine distance and bearing between the new location and the previous location with that of the distance and bearing between the subsequent location and the previous location, to identify cases where birds appeared to move back towards their point of origin shortly after leaving it. In such cases, we assumed that the bird had not moved and deleted the outlier. In practice, it was usually easy to identify such outliers as in many cases not only the immediately preceding and subsequent locations were much closer to each other than either was to the outlier, but so too were many of the preceding and many of the subsequent locations, sometimes during the same transmission cycle. These indicated that the bird was present in much the same area for a considerable length of time and that apparent long-distance short-term movements out of the area prior to final departure were artefacts of poor location accuracy. The speed of travel, estimated as the haversine distance divided by the time between fixes, often confirmed these points as errors, since the speed required to move between the points was unrealistic. As both the spatial and the temporal resolution of fixes generally improved towards the end of the study, it became clear that birds did not undertake long-distance temporary movements from staging, breeding or wintering sites before moving elsewhere, so excluding major outliers in the way described is unlikely to lead to the loss of meaningful information. We assumed that genuine movements had occurred when displacements of >100 km were followed by a series of locations that fell in the same area or along the same bearing. Again, visualising the data in GIS provided confirmation that genuine movements along a biologically meaningful trajectory had taken place. For estimating dates of departure and arrival, we assumed that movements occurred at the mid-point between two successive locations. Departure date from the breeding or wintering grounds was taken as the estimated date on which birds moved 100 km or more (according to the criteria described above) from the area in which they had spent the preceding weeks or months in a direction indicative of migration; in practice, such movements were often very much further than 100 km, or were quickly followed by much longer movements. Arrival date on the breeding grounds was taken as the date on which a bird arrived within the known breeding range of the species in central Kazakhstan, although some birds made subsequent movements to other parts of the breeding range, presumably prospecting for breeding colonies or evading late episodes of severe winter weather.

We clustered the filtered locations by sorting the locations for each tag in chronological order and iteratively calculating the distance from each successive location to the mean centroid of the existing cluster, weighted by location quality (Argos location quality Z carried a weight of 1, location quality 3 a weight of 7, with the intermediate location qualities scored accordingly in intervals of 1). If the location fell within 50 km of the weighted centroid of the current cluster, the location was added to that cluster and the weighted centroid recalculated to incorporate the new point’s coordinates and location quality. If the next location was over 50 km from the centroid, a new cluster was started. “Clusters” containing only one point and those with multiple points but spanning less than 2 days between the first and last locations in that cluster (disregarding the intervals between that cluster and previous and subsequent clusters) were regarded as transit points between places where birds settled, although the very “gappy” nature of the data meant that birds may have spent longer at some of these points than indicated by the clustering protocol and may have spent significant periods at sites for which no locations were received. The length of time bird spent at clusters was estimated as the time between the first and last locations assigned to that cluster plus the sum of half the intervals between that cluster and the previous and the subsequent cluster or transit point. In a small number of clusters (<10%), this yielded estimates of the inferred length of stay that was over twice the time between the first and last recorded locations, because the cluster was preceded or followed by a lengthy absence of records. In these cases we therefore truncated the time the bird was assumed to be present at the site to be no longer than twice the time the bird was known from location timestamps to be present there.

### Colour ringing

Between 2004 and 2015, we fitted 150 adults and 1473 chicks with unique combinations of four colour rings (two on each leg) during the breeding season, most of them in the core study site around the town of Korgalzhyn in central Kazakhstan or along the Irtysh River in Pavlodar province, around 520 km to the north east of Korgalzhyn (see map in Kamp *et al.* 2009). The main aim of colour ringing was to estimate survival rates (Sheldon *et al.* 2013), but we also opportunistically searched for colour-ringed birds during surveys of staging and wintering areas and during intensive breeding season fieldwork at a number of sites across central Kazakhstan.

### Field surveys

Between 2004 and 2017, 88 field surveys were conducted across the range of the Sociable Lapwing (Supplementary Online Fig. S1). They were coordinated by the BirdLife International Sociable Lapwing project and usually implemented on the ground by local conservationists and ornithologists. The surveys lasted between one and three weeks. They aimed at identifying important breeding, stopover and wintering sites, assessing the number of birds that used these sites, identifying local threats and reading colour-rings of bird marked on the breeding ground to track their movements. Areas presumed to hold suitable habitat such as dry steppe areas, semi-desert and wetlands were visited opportunistically and surveyed with telescopes. Searches at the stopover and wintering sites were often guided by satellite-tagged birds and transmitted coordinates were made available to fieldworkers (Donald *et al.* 2016). Birds fitted with satellite transmitters were successfully located in the field in Sudan, Turkey, Kazakhstan and Uzbekistan. All data obtained during the field surveys was incorporated into the sightings database (see below), with a qualifier to allow later filtering.

### Database of historical and recent records

We aimed to synthesize existing data on the distribution of Sociable Lapwing throughout the year and searched a variety of recent and historical sources for records. We systematically searched published English and Russian language literature, such as natural history periodicals, descriptions of local avifaunas from all range states and handbooks, including visits to various libraries and academic institutions in Russia, Kazakhstan and Uzbekistan. Additionally, we screened unpublished reports, again making an effort to include relevant Russian-language publications. Further records, especially from the 19th and early 20th century were obtained from 36 natural history museums across the range (Table S1), where we either visited the collection and copied data from specimen labels, or searched online catalogues. An additional source of records were global and regional e-mail lists over which Sociable Lapwing sighting were shared, as well as websites where birdwatchers and photographers shared information and pictures (Table S2). In all range states, we sent out a request for records to local ornithologists.

Place names were identified using internet gazetteers (including in Russian), Google Earth, Yandex maps and historical maps. Coordinates were extracted for each record and their accuracy classified on a scale ranging from “accurate: coordinates taken with GPS” to “approximate coordinates, accuracy >100 km”. Records extracted from the older literature sometimes used only qualitative measures of abundance, for which we inferred discrete numbers using the protocol shown in Table S3. We excluded records of vagrants from European countries except Ukraine, European Russia and European Kazakhstan as we were interested in distribution and migration patterns across the main range of the species. The completed database contained 3116 observations of Sociable Lapwings, including those from field surveys, made between 1767 and 2019.

### Temperature and land cover data

We assessed the approximate temperature at each satellite location of tagged birds throughout the year by extracting, for each tag location with an Argos location quality of 0-3, the mean monthly temperature of the relevant month, averaged over 1970-2000, at a 5 arc-minute resolution from WorldClim 2.0 (Fick & Hijmans 2017). For analysis of broad land cover types within clusters, we calculated the root mean square of the distance of each point in each cluster from the weighted centroid of that cluster and buffered the weighted centroid by that distance. We then extracted the area of each level-2 IUCN terrestrial habitat class falling within the buffer from Jung *et al.* (2020, submitted) in ArcGIS. We assessed long-term changes in the extent of agricultural land within the species’ range using the HYDE 3.2 historical land use dataset (Klein Goldewijk *et al.* 2017), and shorter-term changes in the extent of irrigated agriculture using the dataset of Siebert *et al.* (2015).

## RESULTS

### Tracking data

Of the 10,364 tracking locations in the filtered dataset, only 2369 (22.8%) were in the higher Argos quality classes of 1, 2 or 3, confirming previous reports that the accuracy of Argos fixes in Central Asia is generally low (Dubinin *et al.* 2010). The temporal resolution of the tags that did not fail quickly was sometimes poor, particularly in the early years of the study, with gaps between successive locations as long as several months, particularly during migration. Applying the filtering and clustering protocol described above reduced the locations to 255 clusters and 248 transit points. In the early years of the project, tag failure was high. We cannot assess whether early failure was due to problems with the tag or harness or due to mortality, but improvements in tag and harness design over the course of the project led to an increase in the longevity of tags so we assume that high rates of failure early in the project were due to tag and harness failure and not to mortality. One colour-ringed bird fitted with a tag that ceased transmission shortly thereafter was relocated the following breeding season alive and apparently well but without the tag or harness. Of the 29 tags fitted, 16 yielded data on one (n = 9) or more (n = 7) complete autumn migrations to the wintering grounds (number of complete autumn migrations tracked = 27), and of these 8 yielded data on one (n = 5) or more (n = 3) complete spring migrations to the breeding grounds (number of complete spring migrations tracked = 13).

### Migration routes

Plotting the filtered satellite tracking data in ArcGIS revealed the existence of two main migration pathways; a longer western route west across Kazakhstan, across or around the western Caspian Sea, then south through the Caucasus and the Levant to wintering areas in Saudi Arabia and eastern Sudan, and a much shorter eastern route due south to Turkmenistan and Uzbekistan, then over or around the mountains of northern Afghanistan to wintering areas in Pakistan and NW India (Fig. 1). The tags or harnesses of five of the 29 tagged birds failed before the migration route could be assessed. Of the remaining 24 birds, 17 took the western flyway and seven took the eastern flyway (exact binomial 95% confidence limits of percentage of birds taking the eastern route: 12.6% - 51.1%). Of the five birds tagged in eastern Kazakhstan, the migratory flyway could be assessed for only two, one of which took the eastern flyway and the other the western flyway. A number of stopover areas were used along both routes (Fig. 1); these are referred to as areas rather than sites because some were very large (birds could stage over 200 km apart in the same stopover area).

**Fig. 1.**
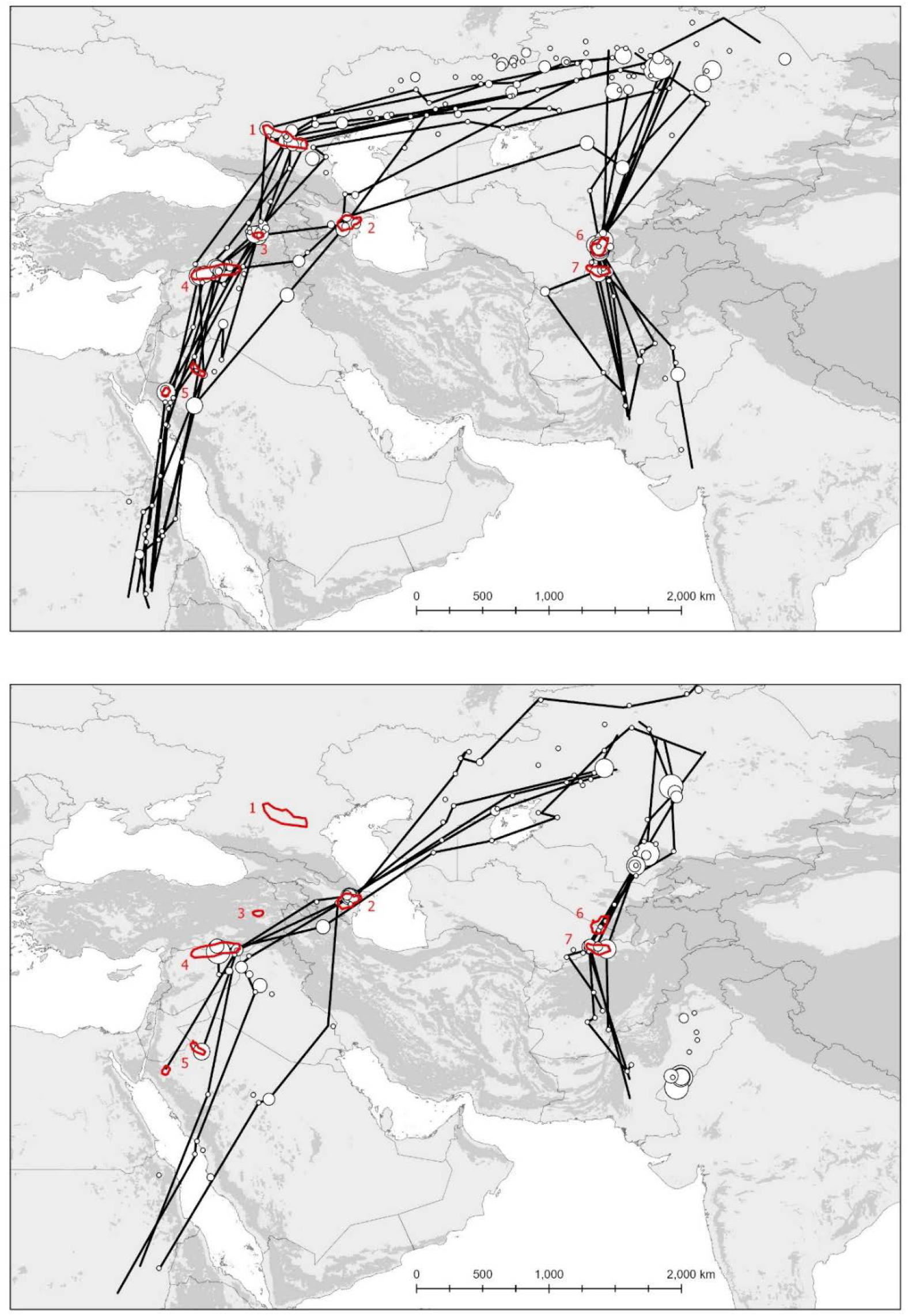
Autumn (upper) and spring (lower) tracks of satellite tagged Sociable Lapwings. Only complete migrations based on data with sufficient temporal resolution to permit confident assessments of the route and stopover locations used are shown. The stopover locations, where birds spent a minimum of two days, are shown as white circles whose relative size represents the amount of time spent at each. Stopover areas that are not part of tracks indicate those used by birds that did not complete a full migration, or for which data were of too low a temporal resolution to permit accurate plotting of the route taken. The tracks include multiple migrations by some individual birds. The main stopover areas used by tagged birds are indicated in red (polygons drawn to include all clusters): 1 = Manych depression, Stavropol Krai, Russia; 2 = lowlands of east-central Azerbaijan; 3 = Malazgirt Plain, eastern Turkey; 4 = irrigated agriculture in southern Turkey and some areas of surviving steppe-like grassland in northern Syria; 5 = pivot-field agriculture in Saudi Arabia (two sites); 6 = Tallymarzhan (Tallymerjen) on the border of Turkmenistan and Uzbekistan; 7 = lowlands of northern Afghanistan. Shaded areas show areas classified as mountains by the UNEP-WCMC Global Map of Mountains (Kapos *et al.* 2000).

The smoothed tracks shown in Fig. 1 averaged 5199 km (range 3998 – 6261 km) on the western flyway and 2839 km (range 2459 – 3547 km) on the eastern flyway. On the western flyway, autumn migration was in most cases though the Stavropol region of south-western Russia and then south through the Caucasus, whereas in spring all tagged birds shortened their return journey to the breeding grounds by crossing the central Caspian Sea directly from Azerbaijan (Fig. 1); neither the stopover area in eastern Turkey nor that in the Stavropol region were visited by any tagged birds in spring, all of which had used one or both in the previous autumn.

On the eastern route, prolonged autumn stopovers were made by all tagged birds at Tallymarzhan and much shorter stopovers at a previously unrecognised stopover area in northern Afghanistan. In spring more use was made of the northern Afghanistan area and birds did not visit Tallymarzhan at all on return migration, or did so only briefly. The stopover area in northern Afghanistan lies less than 100 km south of Tallymarzhan and immediately north of the high mountains at the western end of the Hindu Kush. Birds of the eastern flyway reaching southern Kazakhstan in early spring made often prolonged stopovers at a number of sites, presumably waiting for weather conditions to become suitable for the last movement north to the breeding grounds. In autumn, southward progression was initially faster in birds taking the eastern route, but the prolonged stopover at the Tallymarzhan site on the borders of Turkmenistan and Uzbekistan produced a clear “step” in latitudinal movement, and by the time birds left here they were slightly further north on average than birds taking the southern route (Fig. 2a). In spring, birds on both flyways returned north more rapidly than they had moved south in autumn (Fig. 2a).

**Fig. 2.**
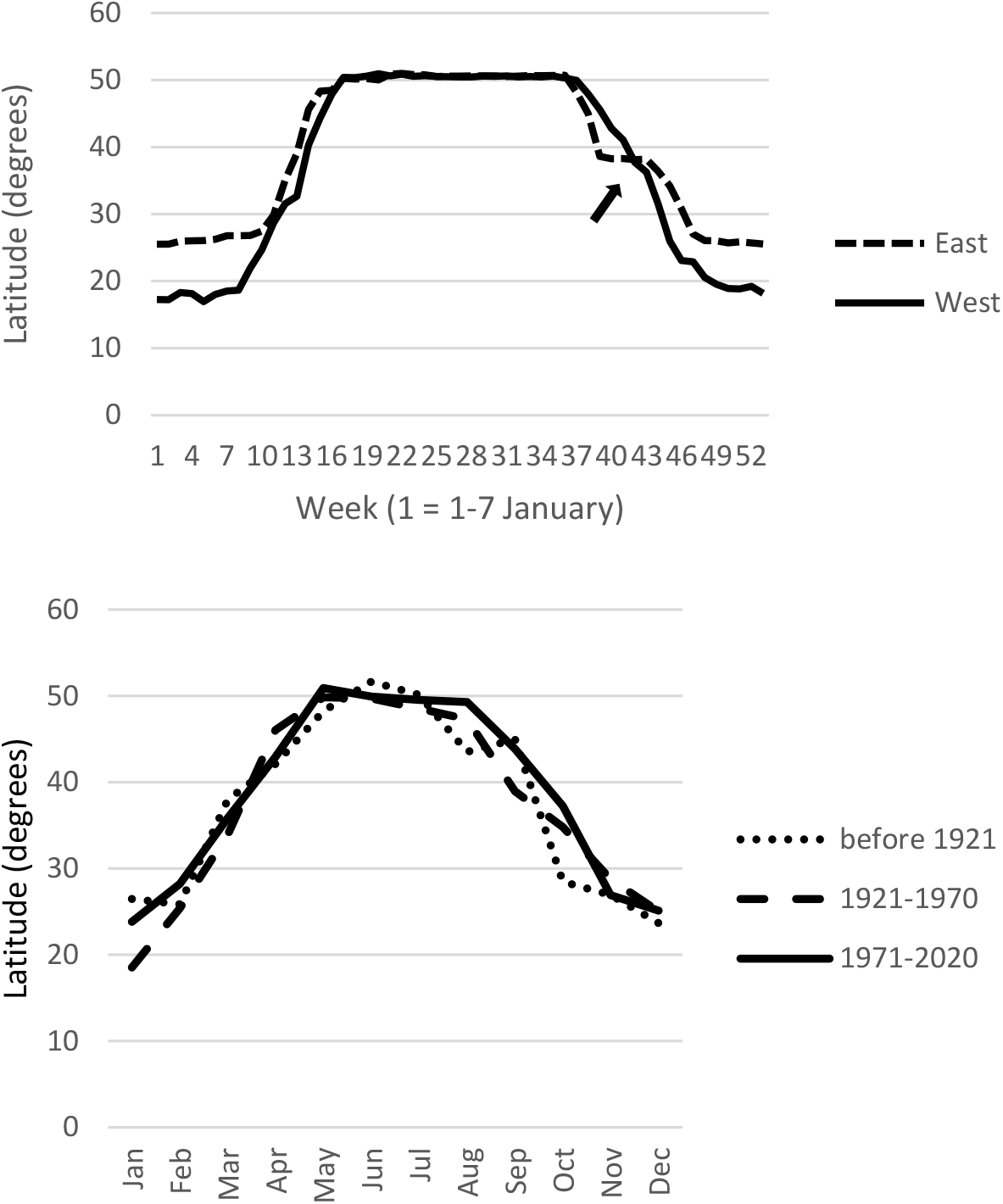
**a.** (upper) Mean weekly latitudinal distribution of satellite-tagged birds of the eastern and western flyways. The ‘plateau’ in the autumn latitudes of eastern flyway birds, indicated by the arrow, represents the lengthy stopover at Tallymarzhan. **b.** (lower) Mean monthly latitudinal distribution of sight records broken down into three time periods; the last 50 years, the previous 50 years and before 1921.

There was a suggestion from the database of field records that median flock size in spring is much smaller than that in autumn (Fig. 3).

**Fig. 3.**
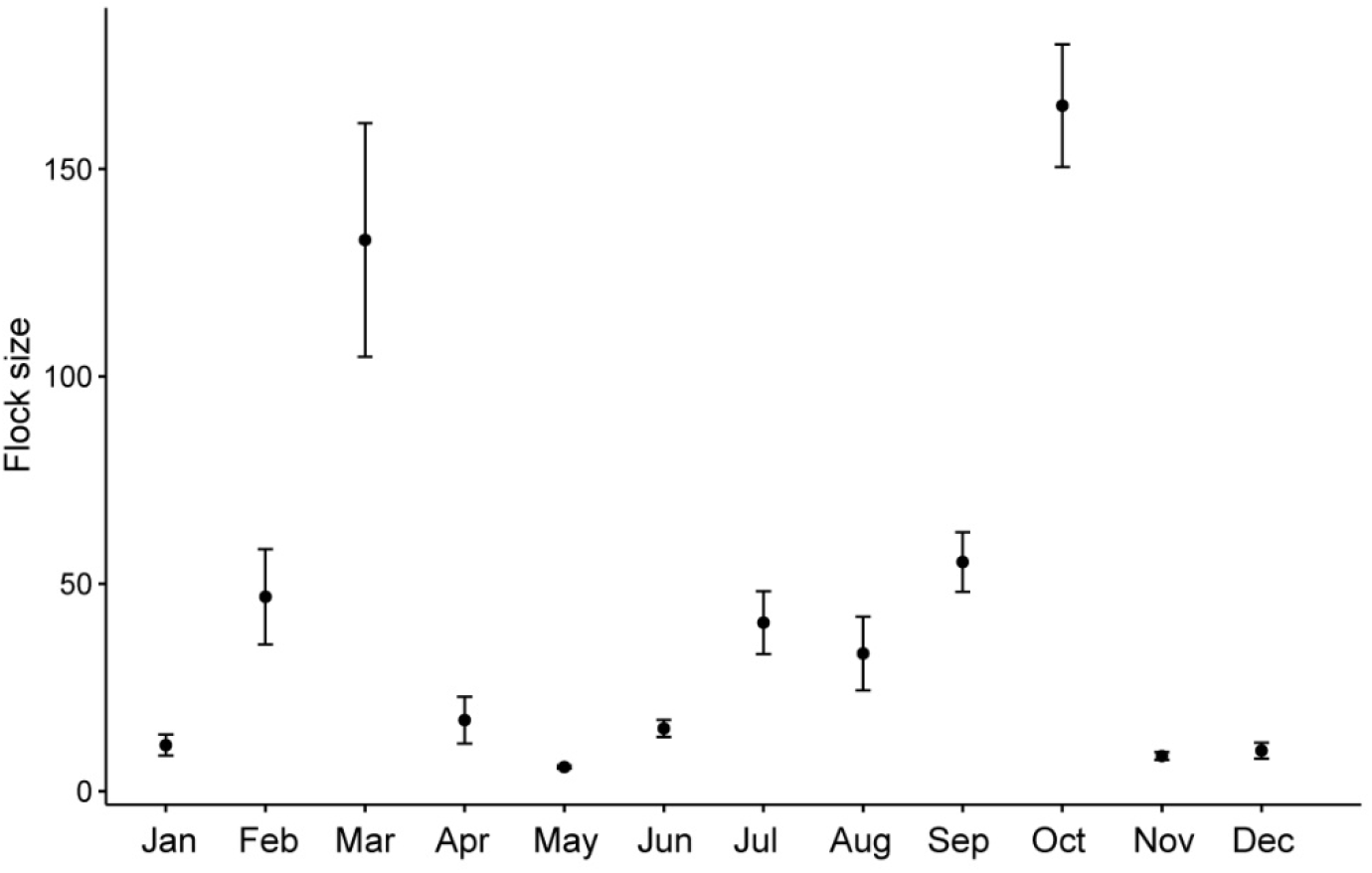
Mean (+/− 1SE) Sociable Lapwing flock size by month 2019 (n= 2286 records from 1970-2019); there were highly significant differences between months in median flock size (Kruskal-Wallis-Test, χ^2^_11_ = 415.93, *P* <0.0001). Flocks were significantly larger during the peak migration months of March and October, compared to the breeding season (May, June) and winter (November to January; pairwise Wilcoxon-tests with Bonferroni correction, *P* <0.001).

Birds on the western flyway wintered in significantly warmer areas than those on the eastern flyway; indeed, temperatures in the wintering areas of western flyway birds were on average higher than those experienced on the breeding grounds in summer (Fig. 4).

**Fig. 4.**
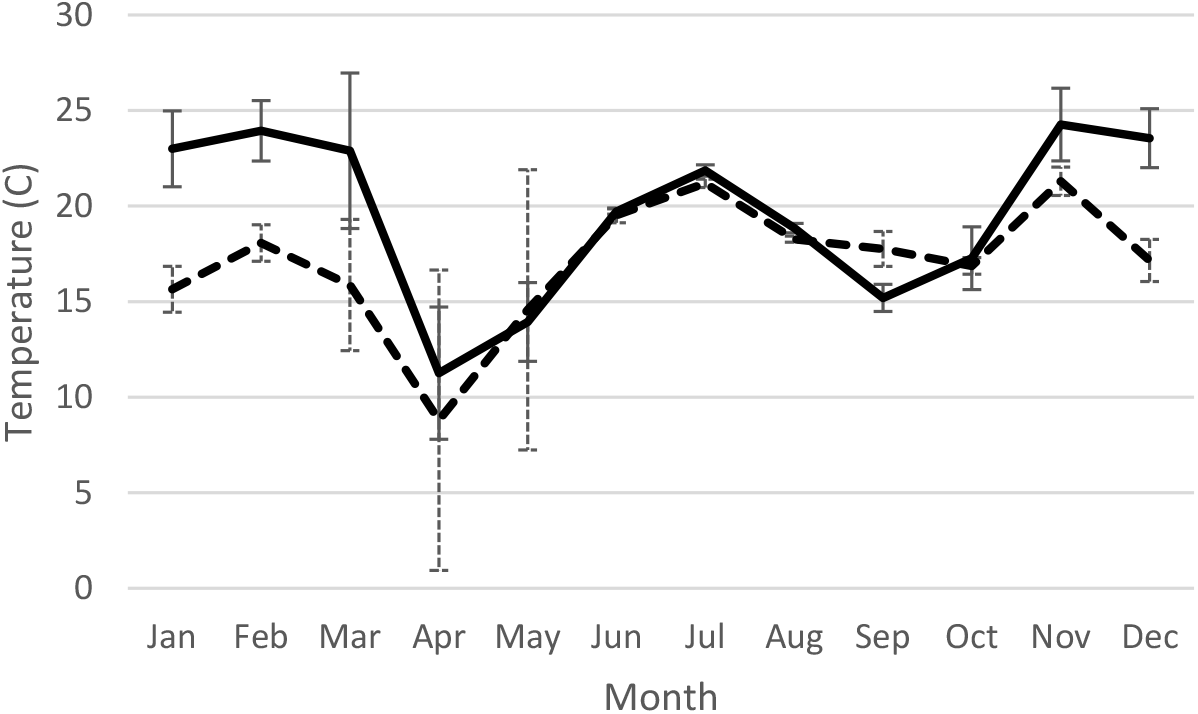
Mean monthly temperature (+/− 1 SE) at point locations of tagged Sociable Lapwings using the eastern (dotted line) and western (solid line) flyways, extracted from BioClim (see Methods). Only locations with Argos quality scores 0-3 were included. Multiple locations of individual birds were averaged by month to avoid pseudreplication.

Birds on both flyways experienced the coldest temperatures of their year in April, when they had just arrived in the breeding grounds or were staging slightly further south, presumably waiting for conditions to improve before moving north again (Fig. 5).

**Fig. 5.**
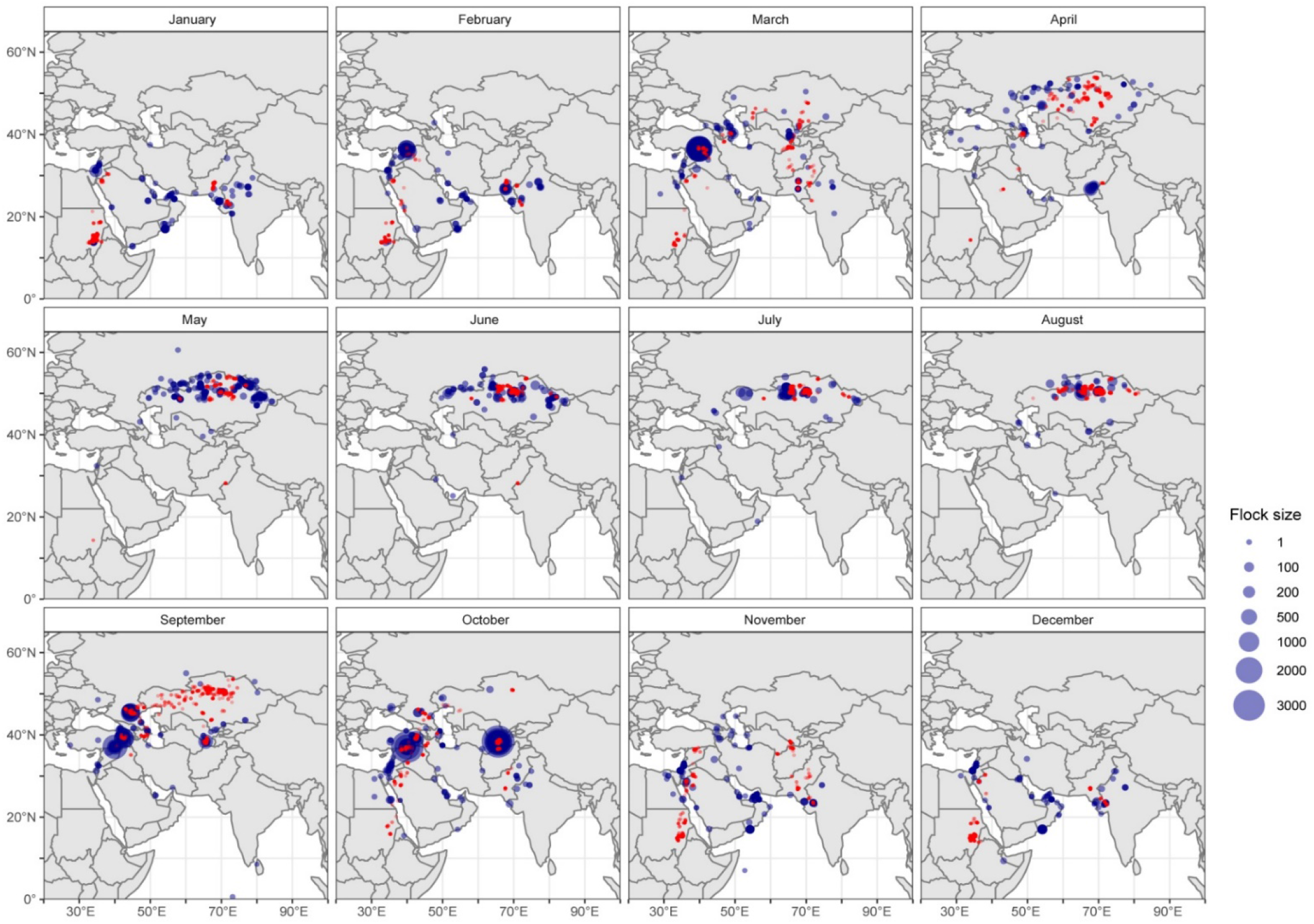
Monthly distribution of satellite tagged birds (red dots) and sight records of the species since 1970 (blue circles, scaled proportional to the number of birds recorded).

Individual tagged birds were consistent in their choice of migration flyway. Of 11 occasions on which birds made second or subsequent migrations (two birds were tracked over four migrations), all took the same route as in the previous year (exact binomial 95% CL of proportion of birds switching flyways: 0 – 0.29), suggesting that if birds ever do move between flyways, it is very unlikely to be a rate greater than one transition per three or four migrations. Birds were also highly consistent in the combination of staging areas visited, and usually visited the same locations within those areas each year. The exception was a single bird that in the autumn of 2014 moved around the north of the Caspian Sea and made a stopover at Manych, but in autumn 2015 moved directly across the Caspian to the staging area in Azerbaijan.

The only other two birds to follow this more direct route in autumn also did so in 2015, although there were also movements by other birds in the same autumn to Manych. Birds visiting staging areas along the western flyway in autumn remained at each for an average of 13.2 days (n = 27 assessments with an accuracy of two days or less, range 3-32), whereas those visiting the single stopover along the eastern route, at Tallymarzhan, remained there an average of 38.1 days (n = 8, range 29-48). In spring, all birds using the western flyway staged in central Azerbaijan but accurate estimates of the length of stay were only available for three birds with stopovers of 5, 10 and 10 days. On the eastern flyway, birds stopped in northern Afghanistan for an average of 7 days (range 5-12) and only one tagged bird stopped at Tallymarzhan, where it remained for 5 days.

Although potentially subject to observer bias in a way that the tagging data do not, the database of historical and recent records of the species picked out the two main migration routes clearly. They also indicated the existence of an additional central flyway, or a branch of the western flyway, not taken by any of our tagged birds, to wintering grounds in coastal regions of the east and south-eastern Arabian peninsula (Fig. 6). This was supported by the recovery of a bird ringed in central Kazakhstan in June 1970, and shot in eastern Iran in October of the same year (Fig. S3). All records of the species in this region have been made in the last 50 years, and this is the only part of the distribution in which there is not also a scattering of older records, suggesting that birds may have started using this region relatively recently in response to the spread of irrigated agriculture there. There was good agreement between the field records and the tagged birds in terms of their temporal distribution, with the field records supporting the tagging data in suggesting a spring migration along the western flyway that largely avoids the stopover areas of the Malazgirt Plain in eastern Turkey and Stavropol in southern Russia, which are used by most birds in autumn (Fig. 5). There was no indication of a long-term change in the latitude at which birds were recorded in each month (Fig. 2b).

**Fig. 6.**
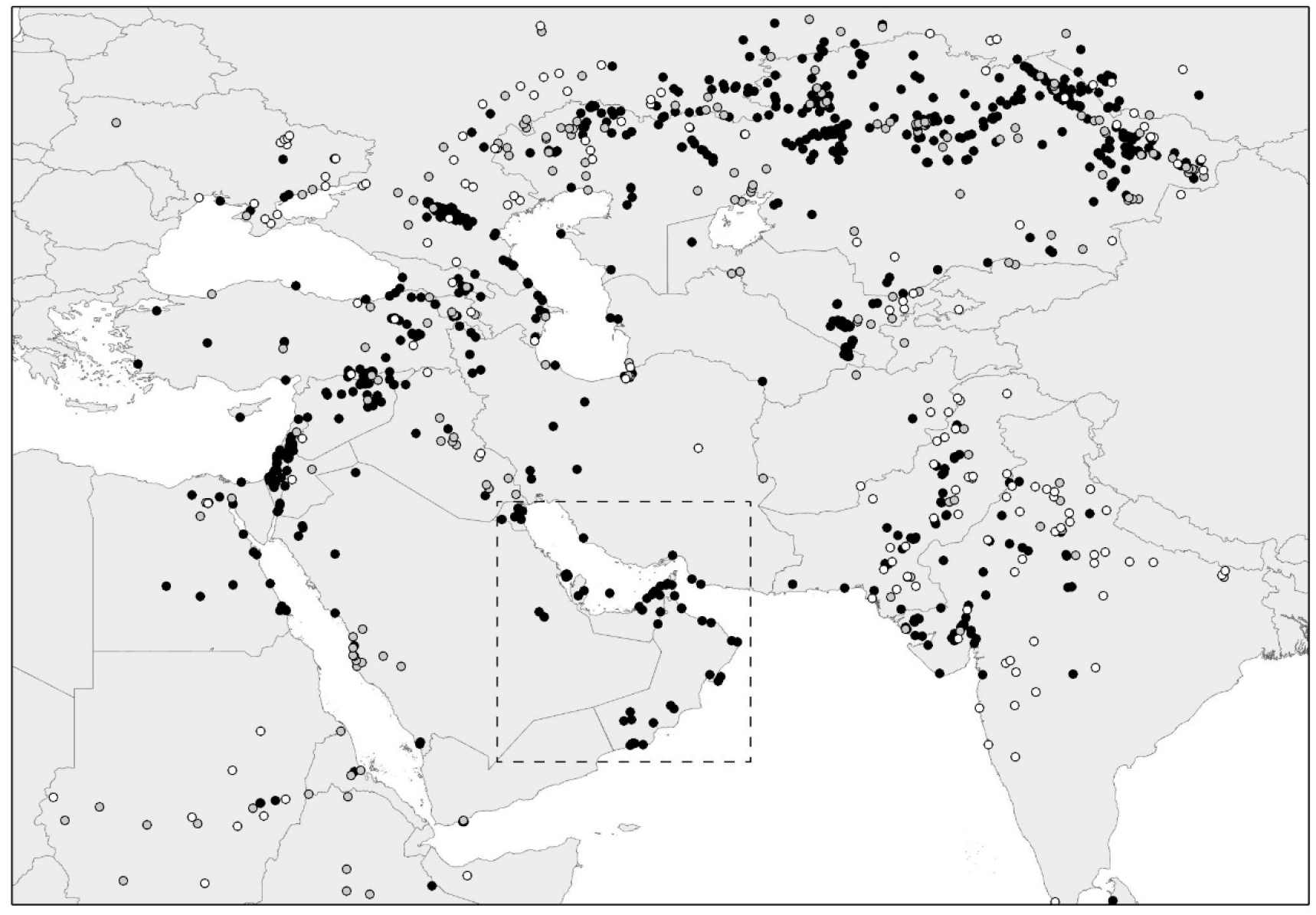
Distribution of sight and specimen records, all months combined. Black dots: last 50 years (1971-2020); grey dots: previous 50 years (1921-1970); white dots: before 1921. All records within the dashed box are from the last 50 years; this is the only region within the species’ range that does not contain older records.

Both flyways required crossing mountain ranges of 2000 m or more, though birds on the western flyway avoided crossing the Greater Caucasus mountains in spring by crossing the Caspian Sea from Azerbaijan (a sea crossing of around 300 km). There was no evidence that birds using the western flyway in autumn avoided crossing the Greater Caucasus Mountains by diverting through the Besh Barmag bottleneck in Azerbaijan, where intensive autumn surveys of migrating birds in autumn in recent years have not recorded significant numbers of Sociable Lapwings (Heiss *et al.* 2020), and all birds following the western route had to cross the Lesser Caucasus and Taurus Mountains. One bird following the eastern flyway showed evidence of avoiding a crossing of the highest parts of the mountains of northern Afghanistan by diverting around their western end (Fig. 1). All birds wintering in Sudan made a crossing of the Red Sea of approximately 300 km.

Only eight Sociable Lapwings colour-ringed as chicks on the breeding grounds in central Kazakhstan were subsequently recorded outside Kazakhstan, five at the Kuma-Manych Depression in Stavropol and three at Tallymarzhan in Uzbekistan (Fig. S3). Logistic regression of a binary variable relating to flyway indicated that there was no significant relationship between choice of migration flyway and latitude or longitude of ringing site or age at ringing (all P > 0.3; Fig. 7). Of two chicks hatched in the same breeding colony less than 2 km apart (although in different years), one was later resighted along the western flyway and one along the eastern flyway.

**Fig. 7.**
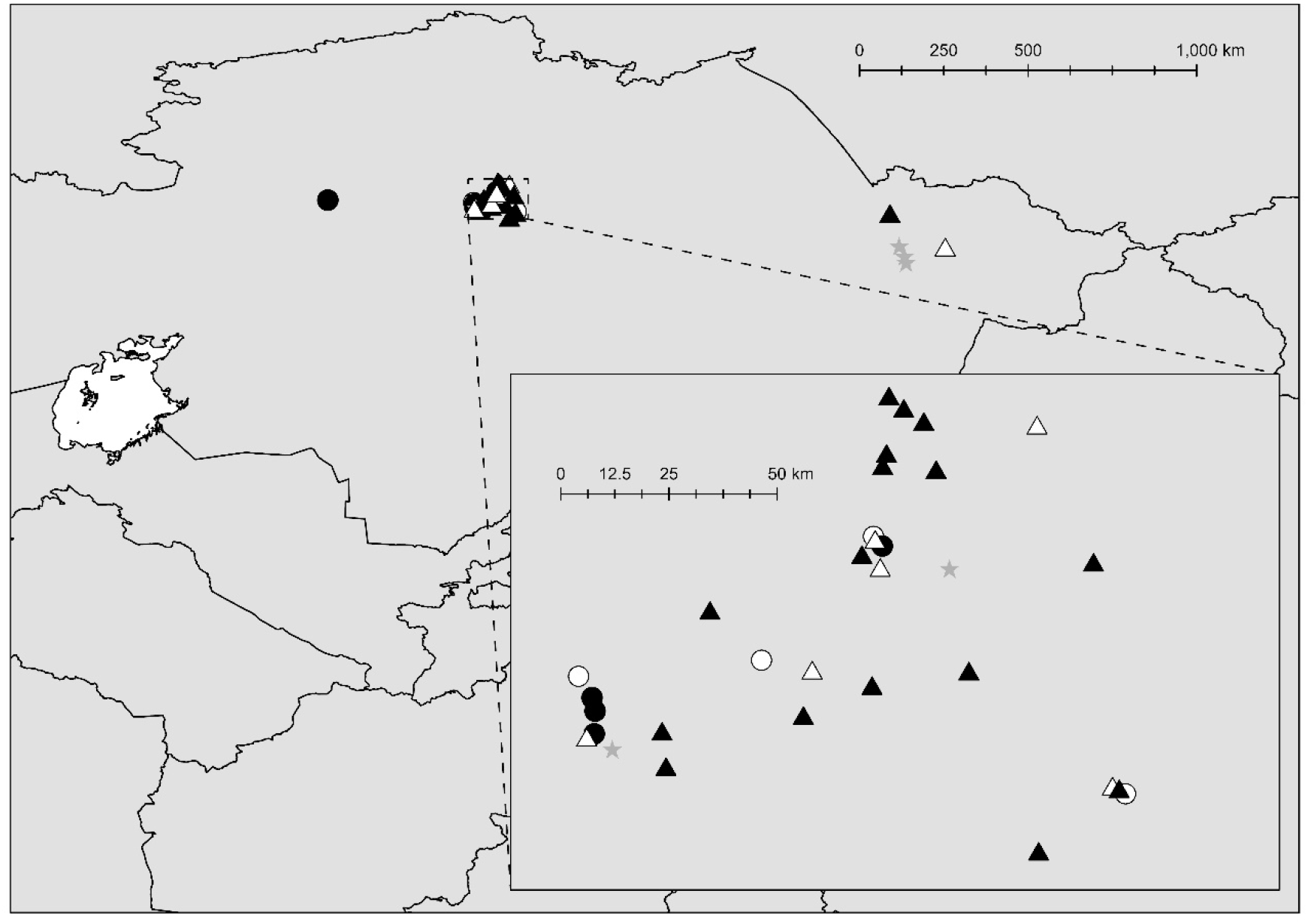
Location of tagging site of adults (triangles) and colour-ringing sites of chicks (circles) that could be assigned to the western (filled symbols) or eastern (open symbols) flyways. The tagging sites of five adults that could not be assigned to a flyway are shown as grey stars.

### Timing and speed of migration

Mean departure and arrival times and mean duration of migration are given in Table 1. There were no significant differences in autumn departure date, arrival date in the final wintering area, spring departure date, return date to the breeding grounds or overall spring or autumn migration period between birds taking the eastern route and birds taking the western route, or between males and females (*t*-tests, all *P* > 0.1). These results were unaffected when seven arrival or departure dates based on intervals between successive locations of 10 or more days were removed. Autumn migration took significantly longer than spring migration (*t*-test *P* < 0.005) and was less synchronous, with a much wider spread of departure and arrival dates (Table 1). On average, Sociable Lapwings spent around a third of the year on the breeding grounds, a third of the year on the wintering grounds and a third of the year moving between them.

**Table 1.**
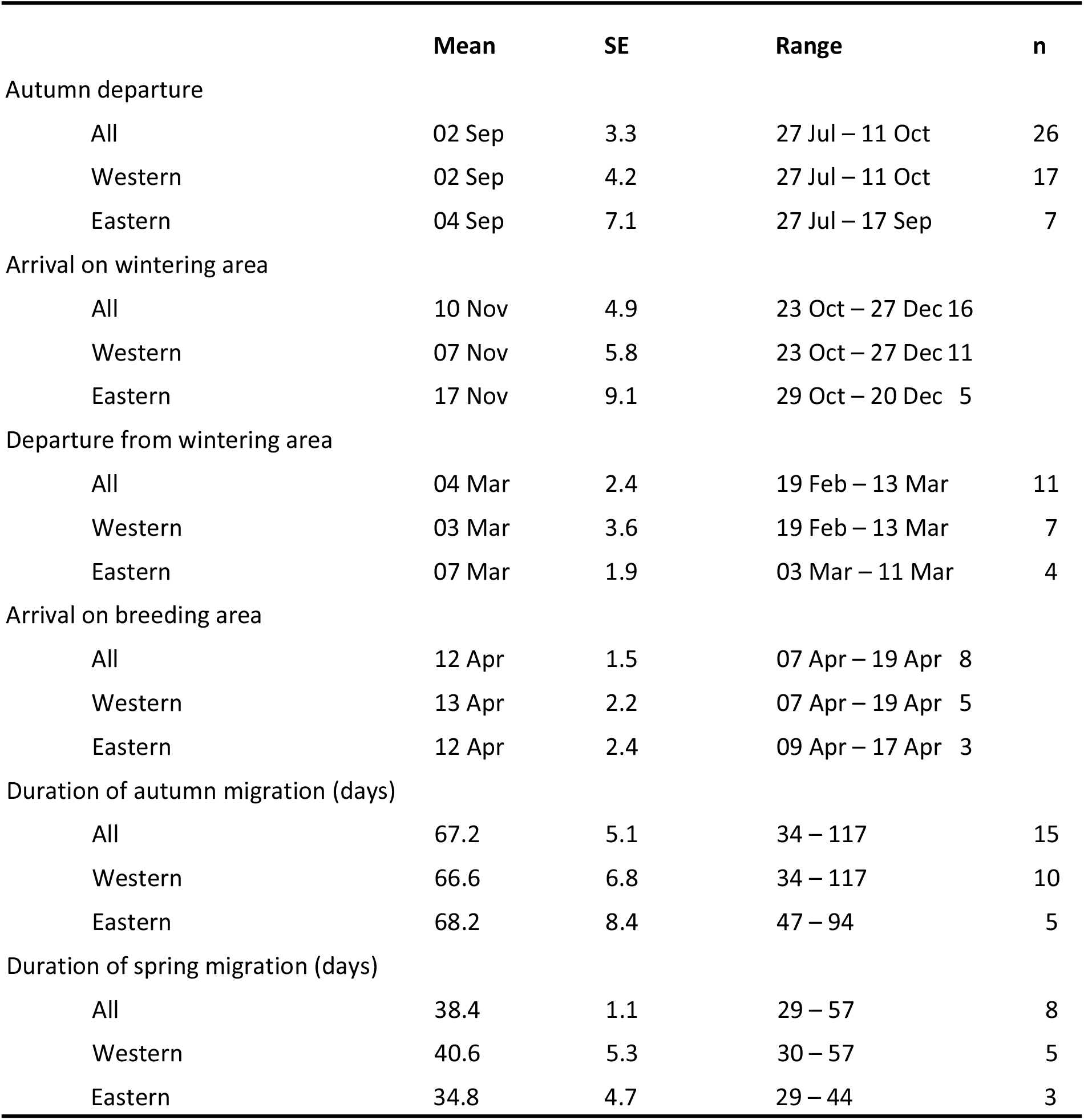
Phenology of migration events of satellite tagged Sociable Lapwings, along western and eastern flyways and both flyways combined. Multiple observations from single birds were averaged over years to reduce pseudoreplication. There were no significant differences between flyways in any of the parameters shown (*t*-tests, *P* > 0.5).

Individual birds were highly consistent in their departure and arrival dates. However, the timing of movements was highly variable between individuals, as shown by the wide range of dates in Table 1. In March and in October, tagged birds were present on both the breeding and the wintering grounds (Fig. 5).

Flight speed during long-distance movements between stopover areas was difficult to estimate due to the generally low temporal resolution of the tracking data, but in six cases where movements of over 1000 km were recorded over an interval between successive transmissions of less than three days, the average was 534 km per day (range 488 – 574). This is likely to be an underestimate of actual flight speeds during long-distance movements since the time spent in flight between successive locations is likely to have been less than the total time between them. The temporal resolution of the tracking data was not sufficient to assess whether birds migrated at night or by day.

### Winter and breeding site fidelity and within-season movements

In winter, birds either remained in the same area (cluster) for the entire period or more frequently moved up to 350 km between areas. The four birds that were tracked over two or more complete winters (two wintering in Sudan and two in Pakistan) were all highly consistent between years in their choice of wintering locations and movements between them (Fig. 8). Three long-distance within-winter movements were recorded; two birds spent part of the winter in Saudi Arabia before moving 1600 km south to Sudan in late December, and one wintered initially in NW India before moving 430 km north into Pakistan in early February. These birds were tracked over a single winter so it is not possible to assess whether this was a pattern they followed each year.

**Fig. 8.**
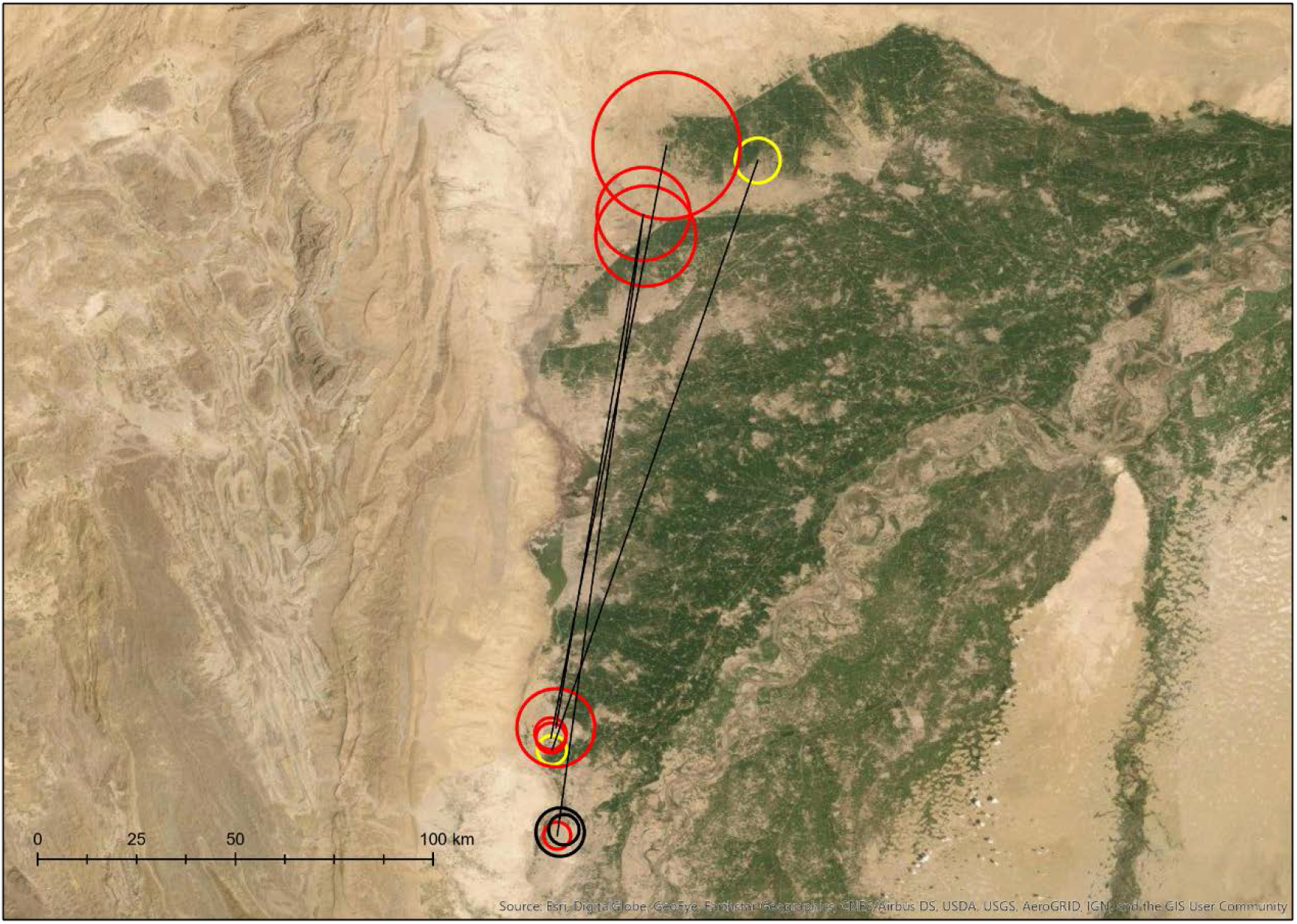
Winter distributions of three tagged Sociable Lapwings in Pakistan. Clusters of records of the three birds are represented by circles proportional in radius to the root mean square distance of records within each cluster from the weighted mean centroid of that cluster. Clusters of records of the same individual within the same winter are connected by lines. One bird (red circles) was tracked over three winters and in each winter used the same two areas. The same two areas were used by another bird (yellow circles) that was tracked over a single winter. A third bird (black circles) was tracked over two winters and used a third area (visited briefly one winter by the first bird), where it spent the entire winter. This shows the consistency of site use by individual birds in winter, and illustrates the selection by birds of areas at the interface of irrigated agriculture and desert.

As in winter, there were within-breeding-season movements of birds between areas that were occupied for several weeks or months, but in contrast to the strong site fidelity shown in winter there was little evidence of strong breeding site fidelity. Of the seven birds that were tracked over two or more summers (including the summer of tagging, which was therefore not a complete season), all three males and three of four females had no spatial overlap whatever between clusters of records in different breeding seasons, and the closest distances between the centroids of any clusters in different breeding seasons were 53 km, 95 km, 305 km for the males and 198 km, 201 km and 291 km for the females.

However, in all these cases comparison was between the year of tagging and the following summer, so it may be that the lack of overlap between years was the product of the incomplete first season or the deterrent effects of capture for tagging (or both). The fourth female was tracked over three complete summers following tagging and showed much greater spatial overlap, with clusters that overlapped across all years. Of the 81 birds ringed as chicks that were re-sighted as adults in subsequent years during the breeding season, the median distance from the natal site was 28.4 km. This is likely greatly to underestimate the degree of natal dispersal since the overwhelming majority of resighting effort was concentrated within a prescribed study area of around 130 km by 120 km. Two birds ringed as chicks in Korgalzhyn were later resighted as adults in Pavlodar, 525 km away. Within the Korgalzhyn region, the mean distance between birds ringed as chicks and later resighted as adults was significantly greater than the distance between birds ringed as adults and subsequently resighted in later years (*t*-test, P = 0.005).

### Habitat use

Across all seasons, 95% of the total area of all clusters comprised just three major land cover types; grassland (47.9%), arable land (35.5%) and desert (11.7%), with negligible areas of wetlands, forest, savanna, shrubland, rocky areas, urban areas and plantations. We therefore used ternary plots to identify differences in the proportions of the three main habitats by season. There was evidence of a greater use of grasslands in the breeding season and of arable land in winter, the plots indicating a selection of areas with intermediate areas of arable land (Fig. 9). With the exception of sites in Saudi Arabia, all the main migration stopover areas and all the areas where birds wintered have had a presence of arable agriculture for at least the last 2000 years (Fig. S2).

**Fig. 9.**
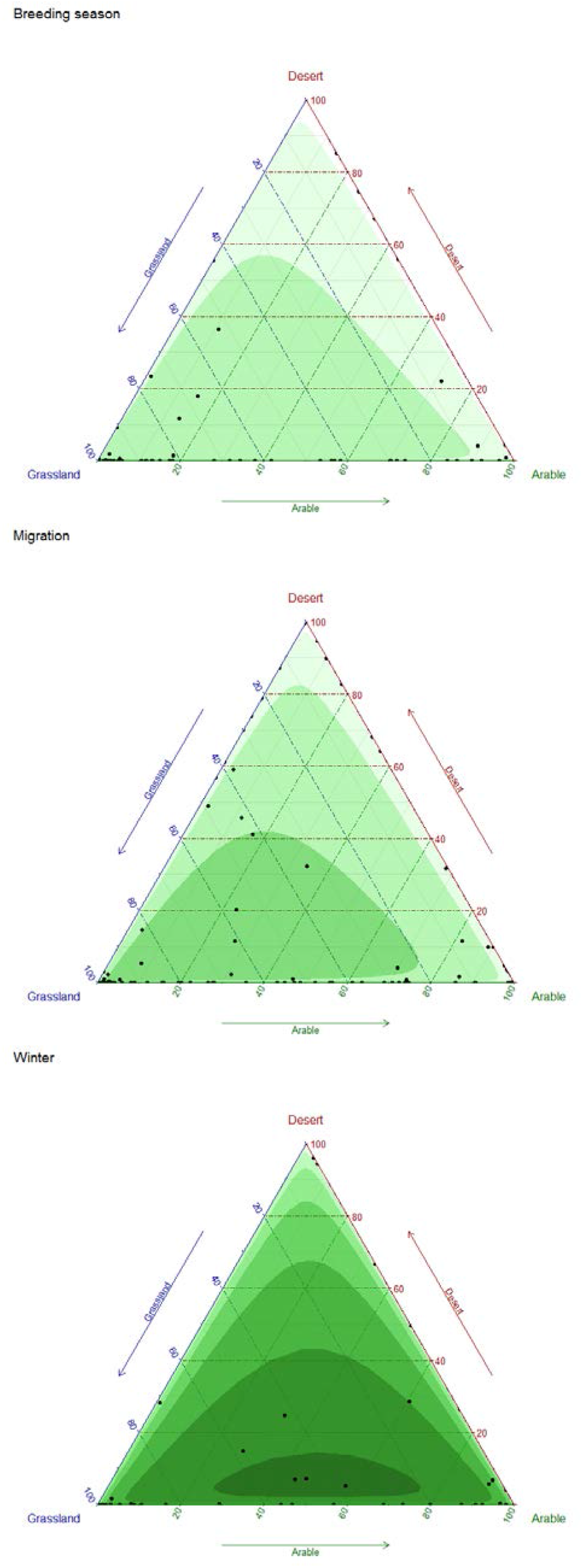
Ternary plots with density contours of the percentage of grassland, arable crops and desert within clusters of records of tagged Sociable Lapwings, broken down by season.

### Survival and population size

Survival is very difficult to estimate from tagging data since mortality events can be difficult or impossible to differentiate from tag or harness failure, particularly when using Argos tags without activity counters (Sergio *et al.* 2019). A staggered-entry Kaplan-Meier plot indicated that the decline over time elapsed since tagging in the proportion of birds known alive approximately exponential (Fig. 10). This allows us to use the Mayfield method and logistic regression to estimate the daily rate of loss to follow-up and to test for differences in this rate between birds on the western and eastern flyways (Aebischer 1999). As expected from inspection of Fig. 11, the daily rate of loss to follow-up was greater for western (0.00285) than eastern (0.00177) flyway birds, but the difference between flyways was not statistically significant (*t* = 1.13, *P* = 0.267). If we assume that all losses to follow-up were deaths, the daily “death rate” based upon pooled data for both flyways was 0.002417, which is equivalent to an annual survival rate of 0.413 (95% confidence limits, 0.290 - 0.588). This is lower than the adult survival rate estimated from resightings of colour-ringed adults (0.66, 95% confidence limits 0.52 - 0.77; Sheldon *et al.* 2012), which is consistent with other information indicating that some loss was due to harness or tag failures.

**Fig. 10.**
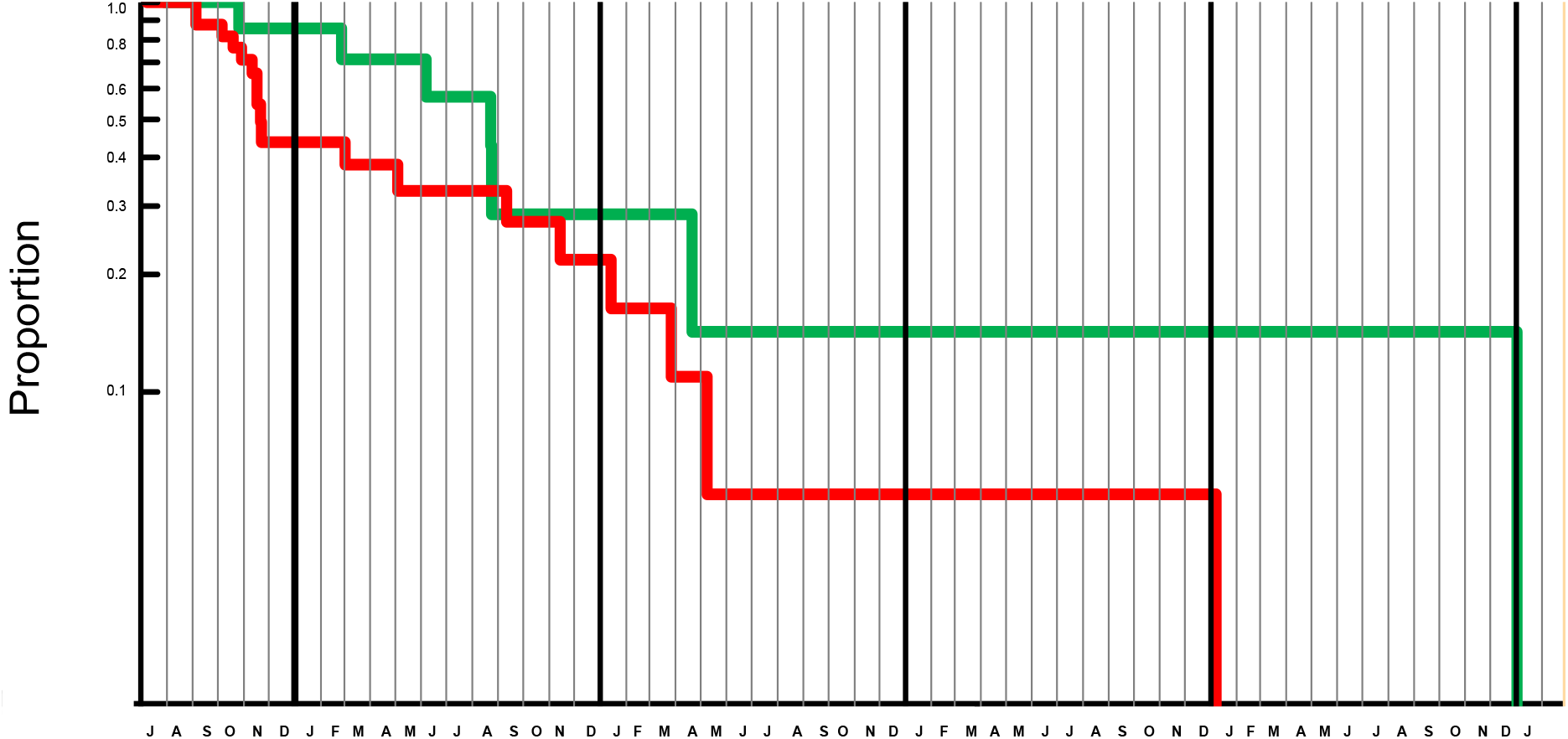
Staggered entry Kaplan-Meier plot showing the proportion of cohorts of satellite-tagged Sociable Lapwings, initially marked in summer in Kazakhstan, remaining alive and carrying functioning tags in relation to time of year. Thin vertical lines show calendar months and thick vertical lines show calendar years. The green line represents birds using the eastern flyway, the red line shows birds using the western flyway. The vertical scale is logarithmic, so if a constant proportion of birds suffered death or tag loss per month, the relationships would be linear. That is approximately the case for the eastern flyway, but there was evidence of higher rates of loss in autumn, winter and spring than in summer on the western flyway.

We attempted to derive a preliminary global population estimate on the two assumptions, both supported by tracking and field survey data, that Tallymarzhan is likely to hold most or all of the eastern flyway population in autumn, and that stopovers were lengthy and hence turnover of birds there during recent counts was low. Tagging data indicated that all tagged birds staged at this site for at least a month and subsequent field visits revealed the presence of large aggregations of birds (Donald *et al.* 2016, Azimov *et al.* 2018). In autumn 2015 the site was estimated by coordinated counts on both sides of the Uzbekistan-Turkmenistan border to hold a total of 6000-8000 birds (Donald *et al.* 2016). Assuming that the proportion of our tagged birds that followed the eastern migration flyway (7 of 24 birds) is representative of the entire population, and taking the exact binomial 95% confidence limits of this proportion (12.6 – 51.1%), the global population can be crudely estimated at (7000/(7/24)) ≈ 24,000 individuals, with rounded 95% CL of 13,700 – 55,560.

## DISCUSSION

The distinctive community of migratory bird species associated with the open steppe habitats and associated wetlands of Central Asia can be divided into three groups on the basis of their wintering areas; those that winter only in Africa (e.g. Black-winged Pratincole *Glareola nordmanni*, Caspian Plover *Charadrius asiaticus*, Red-footed Falcon *Falco vespertinus*, Upcher’s Warbler *Hippolais languida*, Red-tailed Shrike *Lanius phoenicuroides*), those that winter only in southern Asia (e.g. Sykes’s Warbler *Iduna rama*, Paddyfield Warbler *Acrocephalus agricola*, Rosy Starling *Pastor roseus*, Red-headed Bunting *Emberiza bruniceps*) and those that winter in both Africa and southern Asia (e.g. Steppe Eagle *Aquila nipalensis*, Demoiselle Crane *Grus virgo*, Sociable Lapwing, Pallid Harrier *Circus macrourus*, Pallas’s Gull *Ichthyaetus ichthyaetus*, Isabelline Wheatear *Oenanthe isabellina*, Isabelline Shrike *Lanius isabellinus*). The origin of these migratory strategies is unclear, and may be linked to each species’ evolutionary history and its ancestral responses to different conditions in the Plio-Pleistocene (e.g. Bruderer & Salewski 2008) and/or to species-specific requirement for particular environmental conditions that are unevenly distributed across continents (e.g. Somveille *et al.* 2019). There is no indication that the choice of wintering area is linked to the longitude of the centre of the breeding range, and many individuals of species that winter in Africa breed to the east of many individuals of species that winter in Asia. However, in Palearctic species with intercontinental flyways to both Africa and Asia it is usually the case that individuals occupying the western part of the range take the western flyway and those in the eastern part of the breeding range migrate take the eastern flyway, with a clear migratory divide in the breeding range. This pattern has been demonstrated in, for example, Asian Houbara Bustard *Chlamydotis macqueenii* (Combreau *et al.* 2011), a number of tundra-breeding species of the high Arctic (Alerstam & Gudmundsson 1999), European Bee-eater *Merops apiaster* (Hahn *et al.* 2020) and many European-breeding passerines (Møller *et al.* 2011). Red-necked Phalaropes *Phalaropus lobatus* in the western part of the European breeding range migrate westwards to the east Pacific and those in the eastern part of the European breeding range migrate south to the seas off Arabia and East Africa (van Bemmelen *et al.* 2019). We believe that ours is one of very few studies, and perhaps the first, to demonstrate that the choice of intercontinental migration flyway may not be linked to longitude of breeding in adults or longitude of birth in chicks; birds tagged at the western and eastern limits of our sample were equally likely to be part of either flyway, and chicks born even within the same breeding colonies could later select different flyways. A very small sample of Pallid Harriers tagged in central Kazakhstan all wintered in Africa, but a single Pallid Harrier tagged in India also returned to central Kazakhstan (Terraube *et al.* 2012), so the pattern may be common to Central Asian steppe species. Davis *et al.* (2016) tracked Sabine’s Gulls *Xema sabini* from one breeding colony in Arctic Canada, and indeed in one case from one breeding pair in that colony, to both the Pacific and Atlantic oceans and concluded from this that the colony lay on the migratory divide itself. We consider that this is an unlikely explanation for the patterns we observed in Sociable Lapwing, since of the two birds tagged in eastern Kazakhstan, at the eastern edge of the species’ breeding range, one took the western flyway and one the eastern.

In other respects, the spatial and temporal patterns we documented in Sociable Lapwings reflect those noted in other shorebird species; for example, in the faster spring than autumn migration (e.g. Duijns *et al.* 2019), and in the high individual consistency but considerable between-individual variation in the timing and direction of migration and in the use of staging sites along the migration routes (Battley 2006, Gill *et al.* 2019, Méndez *et al.* 2020). However, our results suggest that while birds were highly faithful to their migration stopover areas and wintering areas, they appeared to show much lower fidelity to breeding sites. The reasons for this are unclear, but may be related to the species’ requirement for very short-grazed steppe for breeding, which in natural steppe systems is produced by herds of native migratory ungulates and by fire, both of which may be spatially unpredictable (Kamp *et al.* 2009, Dara *et al.* 2019). Thus a strategy of strict breeding site fidelity may be less adaptive than in species in less variable habitats.

Sociable Lapwing is one of a number of globally threatened birds that rely largely or wholly on agriculture (Wright *et al.* 2011). During migration and on the wintering grounds, the species appears to be strongly associated with areas of agriculture, particularly along rivers.

Several of the main stopover sites identified by this study have already been studied in greater detail (Field *et al.* 2007, Hofland & Keijl 2008, Biricik 2009, Kashkarov *et al.* 2012, Asswad 2014, Donald *et al.* 2016), and these largely confirm the use made by birds of arable agriculture except in northern Syria and at Tallymarzhan, where some surviving areas of steppe-like habitats were used. The main staging areas are in areas along the Manych, Euphrates and Amy Darya (the Oxus of ancient times) and the main wintering areas along the Nile and Indus systems. These areas are home to some of the world’s most ancient agricultural systems, and the migratory and wintering distributions of the Sociable Lapwing may reflect an association with crop cultivation that goes back thousands of years. More recently, the appearance of irrigated agriculture in the Arabian Peninsula, particularly in the form of centre-pivot irrigation that brings up water from fossil aquifers, appears likely to have changed the migratory patterns of the species, allowing birds to stage or even winter in areas that would previously have been unsuitable (Babbington & Roberts 2017). This form of agriculture, which first appeared in the 1950s but expanded greatly after the 1980s, has had generally beneficial impacts on a number of bird species (Campbell & Smiles 2019), and has allowed Sociable Lapwings to shorten their migratory pathways and perhaps to establish new ones. Although highly biased by the distribution of observers, the distribution of sight records of the species suggests that the use of the Gulf regions of the Arabian Peninsula, where birds are always found in close proximity to irrigated crops, may be a recent phenomenon. Steppe Eagle may similarly have benefitted from human activities in the Arabian Peninsula, such as farming and the creation of open refuse and offal disposal tips, and large number of birds have started to winter there; indeed the largest aggregation of the species ever recorded was discovered in two waste disposal areas in the winter of 2019-2020 (Keijmel *et al.* 2020). The existence of a small central flyway of Sociable Lapwings is supported by the regular presence in winter of a small population of Sociable Lapwings and from the few records from Iran, which peak in November (Ashoori *et al.* 2013), by which time birds on the main flyways are already on their wintering grounds. There may also be a very small wintering population in coastal Iran, which again appears to be a new wintering area (Ashoori *et al.* 2013). None of the 23 tagged birds for which the final wintering destination could be determined (one of the 24 birds which it was possible to assign to the western flyway failed before its final wintering destination could be assessed) used this putative third flyway. If we assume that we tagged a sample representative of the population as a whole, the maximum proportion of birds using this flyway is likely to be small (exact binomial upper 95% confidence limit = 14.8%). Other flyways may also exist; the regular wintering of small numbers of birds in Iberia, and their apparently high survival there, may represent the relic of a far-western flyway from times when the species’ distribution extended much further west (de Juana 2011).

Sociable Lapwings are widely, thinly and unpredictably dispersed over huge and often inaccessible areas on both the breeding and wintering grounds, making estimation of the global population size of this species extremely difficult. Their concentration in a small number of predictable staging areas when on migration offers the best opportunity to gather information on population size and trends. Our estimate of a global population of 23,400 individuals (rounded 95% CL: 13,700 – 55,560), based on counts at one such site and an estimate of the proportion of the entire population using that site, is not greatly different from the most recent estimate of 16-17,000 individuals, estimated from extrapolations of densities on the breeding grounds (BirdLife International 2020), and the 95% CL of our estimate encompass that estimate. Further coordinated cross-border counts at Tallymarzhan offer the best opportunity to track changes in the eastern flyway population, but these may not reflect trends in the western flyway population. Birds using the western flyway in autumn did not use all the available stopover areas along that flyway and were often present for fairly short periods at the sites they did use, indicating a high turnover of birds using these sites; therefore counts of birds at autumn stopover areas would be unlikely to yield accurate estimates of the size of the western flyway population. Estimating trends in the western flyway population is further complicated by the sometimes large size of the stopover areas (the autumn staging areas at Manych and Ceylanpinar can hold birds staging over 200 km apart) and by the poor security situation in some range states, incl. the large stopover sites in the Turkish-Syrian border region. Wintering and staging areas in Saudi Arabia are smaller and more easily accessed but not all birds use them, and those that do often spend less than a week there, so turnover is likely to be high. Our results suggest that the best opportunity to track trends in the western population may be to undertake coordinated counts in Azerbaijan during spring migration, where all tagged birds using the western flyway staged for between 5 and 10 days. While some field surveys have been undertaken in Azerbaijan in autumn (Vidal & Sheldon 2016), when most tagged birds did not pass through that country, no systematic searches for the species have yet been undertaken there during spring migration.

Hunting of Sociable Lapwings along the western migration flyway, particularly in Syria and Iraq, has been identified as a key threat to the species (Sheldon *et al.* 2012), and is supported by photographic evidence (Fig. S4). According to the World Database on Protected Areas (WDPA: https://www.protectedplanet.net/), none of the stopover sites identified in Fig. 1 has any coverage by protected areas (though Turkey and Syria do not report their protected areas to the WDPA). Recent estimates of birds illegally killed in countries along the Mediterranean (Brochet *et al.* 2016), the Caucasus (Brochet *et al.* 2019b) and the Arabian Peninsula, Iran and Iraq (Brochet *et al.* 2019a), suggested that a minimum of 76-630 Sociable Lapwings are killed annually. Hunting of wild birds is particularly severe in the Caucasus region, through which all western flyway Sociable Lapwings pass in autumn, and is highest in Azerbaijan (Brochet *et al.* 2019b), which may hold the entire flyway population in spring. This makes the need for spring surveys of the species in Azerbaijan particularly urgent. No data on the hunting of Sociable Lapwings exist for the countries of the eastern flyway but it is unlikely to be on the scale of that on the western flyway. No such threats have been recorded at the species’ main migration stopover at Tallymarzhan, where the birds’ approachability suggests that they are not hunted (Donald *et al.* 2016, Azimov *et al.* 2018). Low adult survival has been identified as the most likely demographic driver of decline (Sheldon *et al.* 2013). In the absence of known threats to adult survival on the breeding grounds, and with the information presented here on the species’ selection of agricultural habitats that are at least stable in area and in many places are increasing, we suggest that illegal hunting of birds along the western flyway in spring and autumn is, on current evidence, the most plausible cause of recent population declines.

## Supporting information

Donald et al._suppl material

## ACKNOWLEDGEMENTS

This study was partly funded by grants from the Darwin Initiative of the UK Government (grant references 15-032, 18-004 and EIDPO035). Additional funding was provided by Swarovski Optik (the BirdLife Species Champion for Sociable Lapwing) through the BirdLife Preventing Extinctions Programme, the African-Eurasian Waterbird Agreement (AEWA) and the German Ornithological Society (DO-G), the Mohammed bin Zayed Species Conservation Fund (MBZ) and the Ornithological Society of the Middle East, the Caucasus and Central Asia (OSME). We thank the many people for sharing their unpublished Sociable Lapwing records, often accompanied by detailed notes and photographs, and the many surveyors who took part in field surveys: they are listed in Appendix S1. Ralf Aumüller, Nigel Collar, Andrey Kovalenko, Anna Ten, Pavel Tomkovich and Christian Wegst visited various museums and copied information from the labels of Sociable Lapwing specimens. We also thank the many students from the Universities of Karaganda, Petropavlovsk, Kostanai, Nur-Sultan (Astana), Almaty and Tashkent who joined the field teams as part of a parallel programme of training.

## Data availability statement

The data that support the findings of this study are available on request from the corresponding author. The data are not publicly available due to privacy or ethical restrictions.

## SUPPORTING INFORMATION

Additional Supporting Information may be found in the online version of this article.

**Fig. S1.** Geographical and seasonal distribution of targeted field surveys for Sociable Lapwings between 2004 and 2017.

**Fig. S2.** Estimated distribution of croplands in range of Sociable Lapwing c. 2000 years before present.

**Fig. S3.** Recoveries of birds ringed on the breeding grounds.

**Fig. S4.** Illegally hunted Sociable Lapwings, probably Syria.

**Table S1.** Museums surveyed for Sociable Lapwing specimens and clutches and number of records obtained.

**Table S2.** Online resources regularly checked for Sociable Lapwing records between 2005 and 2020.

**Table S3.** Conversion of qualitative descriptions of numbers of birds into numeric estimates.

**Appendix S1.** Extended Acknowledgements

