## Supplementary material for "Migration strategy and site fidelity of the globally threatened Sociable Lapwing *Vanellus gregarius*": Donald et al._suppl material

Fig.  S1. Geographical and seasonal distribution of targeted field surveys for Sociable Lapwings between 2004 and 2017.

Fig. S2. Estimated distribution of croplands c. 2000 years before present, from the HYDE database. All the currently used migration stopover areas (in red) and all the sites used by satellite tagged birds in winter (black circles) are in areas that have been cropped for at least two millennia.

Fig. S3. Recoveries of birds ringed on the breeding grounds in 1970 (1 bird with metal ring, shot) and 2004–2015. Only recoveries of at least 70 km distance the ringing site are plotted. In panel A, all records from the sightings database are plotted in the background (black dots), in panel B, all satellite tracking fixes available for analysis. Note that all but one recoveries of colour-ringed birds were made during targeted surveys at stopover sites.

Fig. S4. Illegally hunted Sociable Lapwings displayed by hunters, probably Syria. Anon.

Table S1. Museums surveyed for Sociable Lapwing specimens and clutches and number of records obtained. Given is the number of birds/clutches for which at least the year in which and the site where they were shot or collected was marked on the collection voucher.

Table S2. Online resources regularly checked for Sociable Lapwing records between 2005 and 2020, or between 2005 and the year the platform was discontinued.

Table S3. Conversion of qualitative descriptions of numbers of birds into numeric estimates.

Appendix S1: Extended Acknowledgements

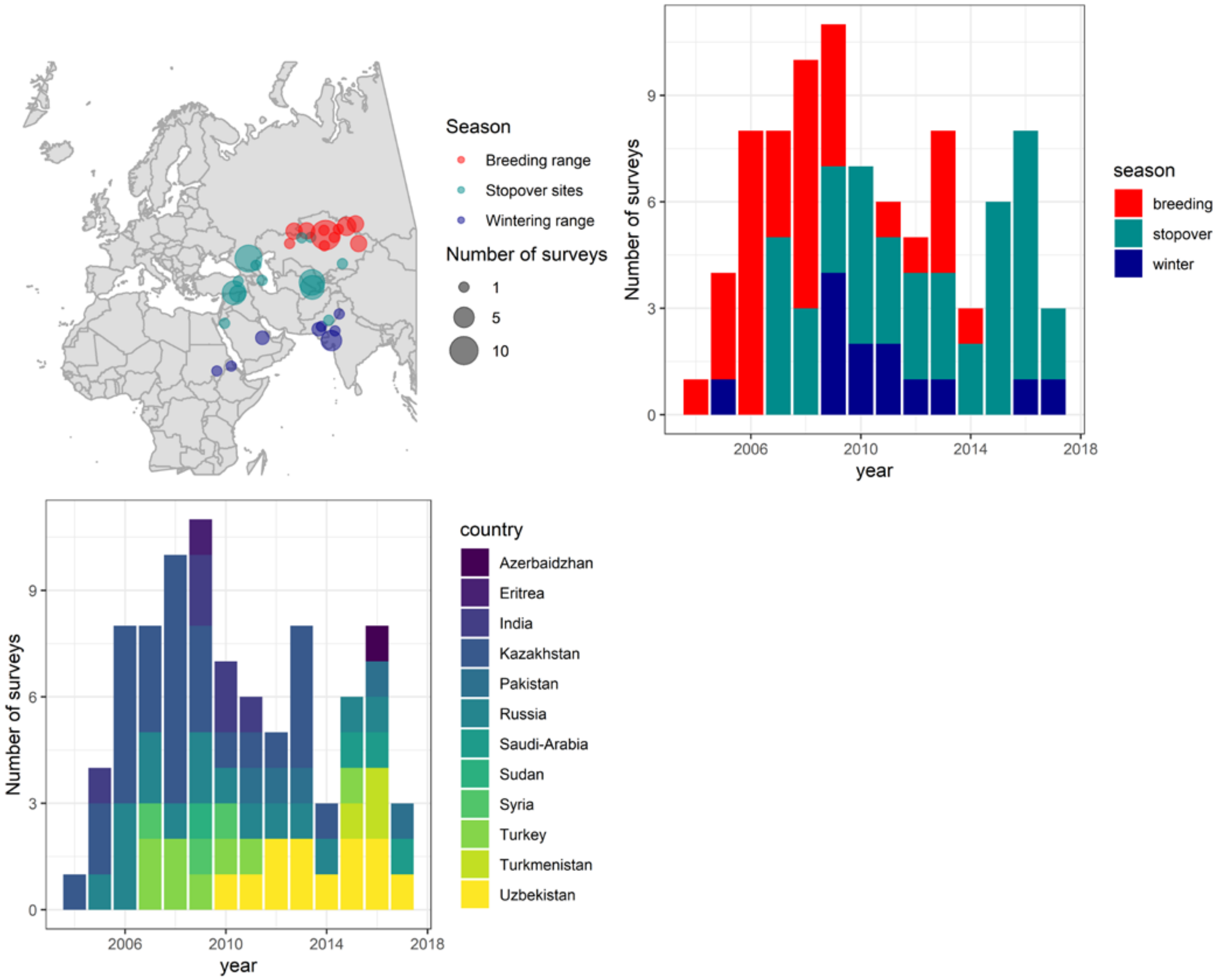


**Fig. S1.** Geographical and seasonal distribution of targeted surveys for Sociable Lapwings between 2004 and 2017.

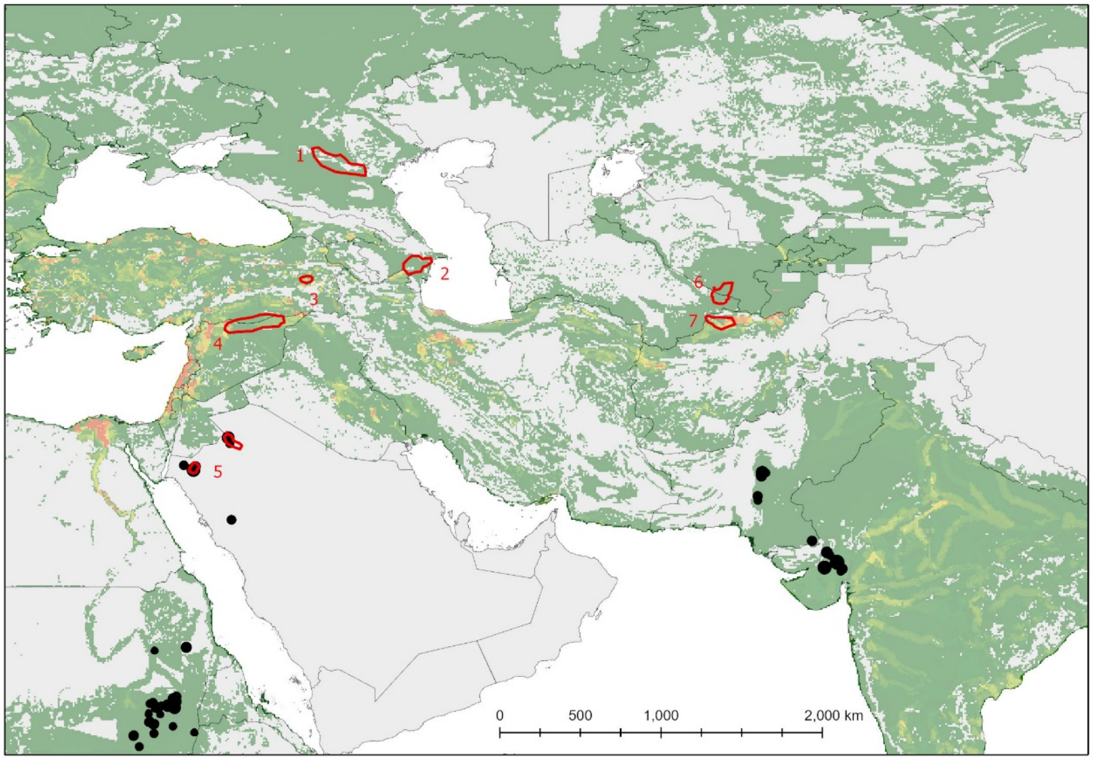


**Fig. S2.** Estimated distribution of croplands c. 2000 years before present, from the HYDE database. All the currently used migration stopover areas (in red) and all the sites used by satellite tagged birds in winter (black circles) are in areas that have been cropped for at least two millennia.

**Fig. S3.** Recoveries of birds ringed on the breeding grounds in 1970 (1 bird with metal ring, shot) and 2004–2015. Only recoveries of at least 70 km distance the ringing site are plotted. In panel A, all records from the sightings database are plotted in the background (black dots), in panel B, all satellite tracking fixes available for analysis. Note that all but one recoveries of colour-ringed birds were made during targeted surveys at stopover sites.

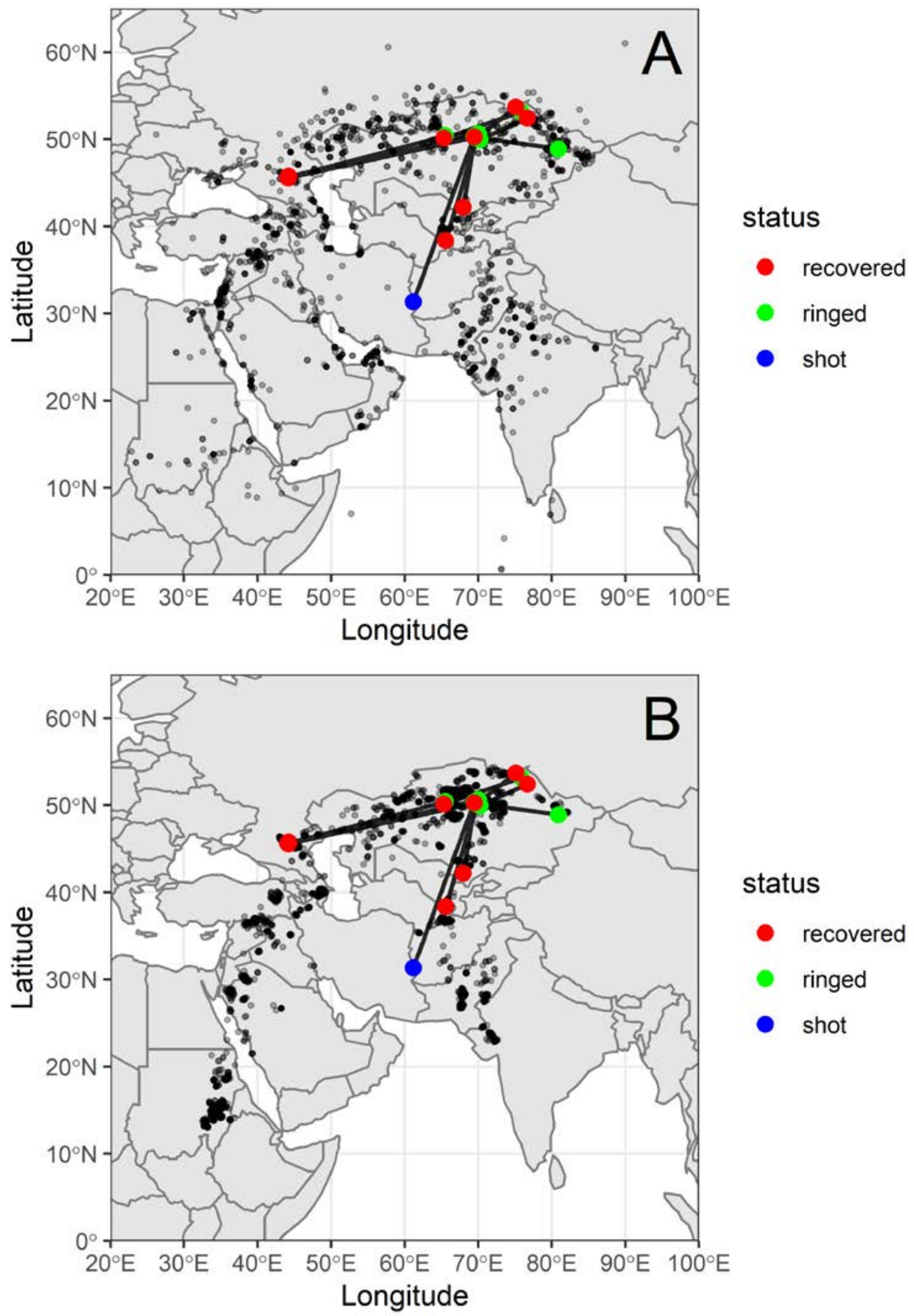

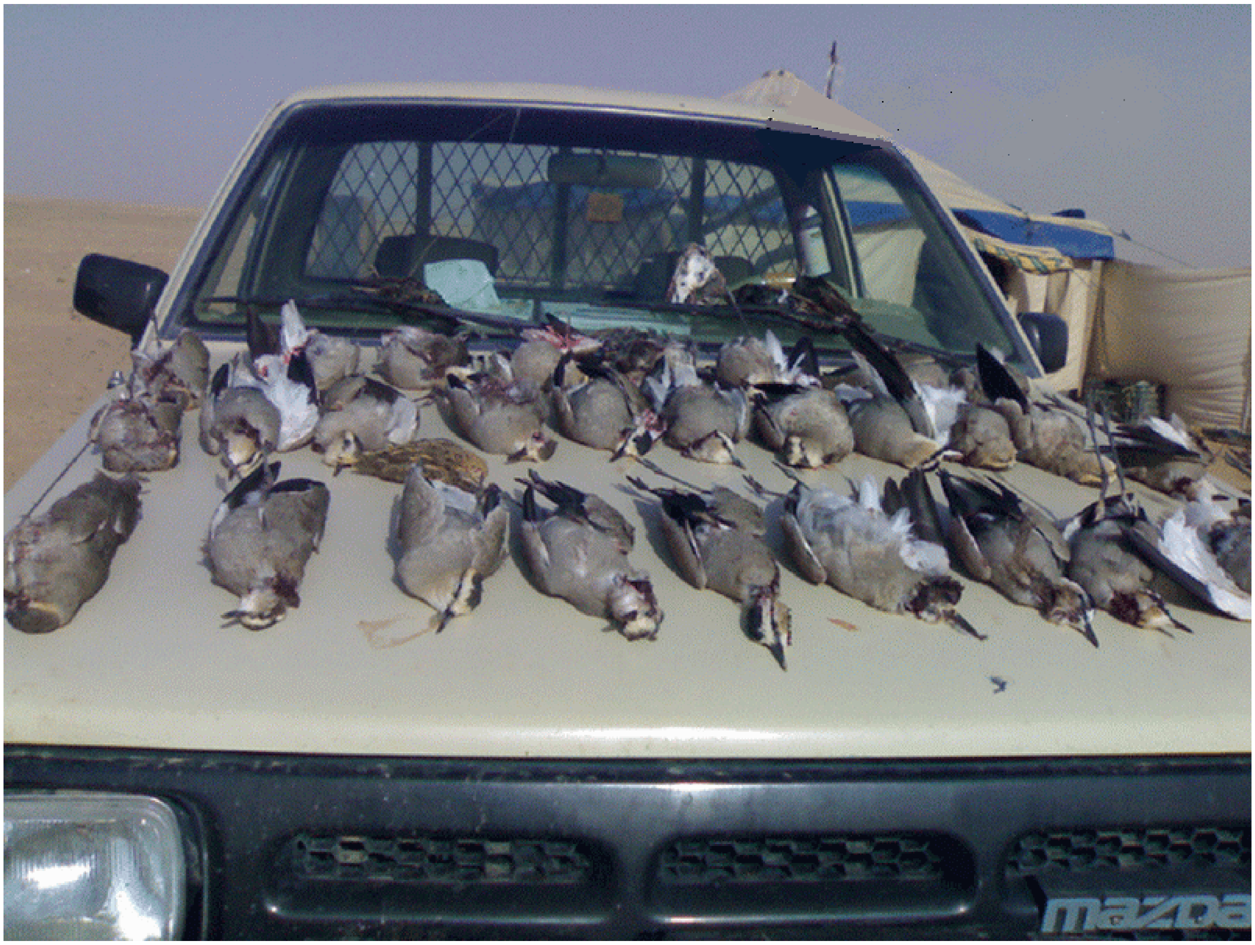


Fig. S4. Illegally hunted Sociable Lapwings displayed by hunters, probably Syria. Anon.

**Table S1.** Museums surveyed for Sociable Lapwing specimens and clutches and number of records obtained. Given is the number of birds/clutches for which at least the year in which and the site where they were shot or collected was marked on the collection voucher.

| Museum | Country | Number of specimens | Number of clutches | Online catalogue |
|---|---|---|---|---|
| Institut Royal des Sciences Naturelles de Belgique | Belgium | 4 | | 1 |
| University of Ghent, Zoology Museum | Belgium | 1 | | 1 |
| Musée Zoologique de la Ville de Strasbourg | France | 2 | | 1 |
| Museum für Naturkunde Berlin | Germany | 4 | | |
| Oldenburg Natural History Museum | Germany | 1 | | 1 |
| Rosensteinmuseum Stuttgart | Germany | 2 | | |
| Zoologische Staatssammlung München, museum specimen | Germany | 2 | | 1 |
| Bombay Natural History Museum | India | 6 | | |
| Collection of the Insitute of Zoology, Almaty | Kazakhstan | 98 | | |
| Rijksmuseum van Natuurlijke Histoire, Leiden | Netherlands | 5 | | |
| Zoological Museum Amsterdam, museum specimen | Netherlands | 4 | | 1 |
| Kazan university museum, Tatarstan | Russia | 1 | | 1 |
| Khvalinsk local history museum, Saratov oblast' | Russia | 1 | | 1 |
| Moscow state university museum | Russia | 65 | | |
| Vol'zhsk local history museum, Saratov oblast' | Russia | 1 | | 1 |
| Zoological collection of the Pedagogic institute, Saratov state university | Russia | 2 | | 1 |
| Museu de Ciènces Naturals de Barcelona | Spain | 1 | | 1 |
| Museum of Evolution, Uppsala, Sweden | Sweden | 1 | | 1 |
| Natural History Museum, Tring | UK | 93 | 18 | |
| University Museum of Zoology Cambridge, Bird collection | UK | 3 | | |
| Museum of the Kharkov National University | Ukraine | 8 | | 1 |
| Academy of Natural Sciences, Philadelphia (AMSP) | USA | 2 | | 1 |
| Academy of Natural Sciences, Philadelphia (AMSP) | USA | 2 | | 1 |
| American Museum of Natural History (AMNH), New York | USA | 2 | | 1 |
| California Academy of Sciences (CAS) | USA | 1 | | 1 |
| Cornell University Museum, New York | USA | 1 | | 1 |
| Field Museum of Natural History, Chicago (FMNH) | USA | 17 | 2 | 1 |
| Museum of Comparative Zoology, Harvard University | USA | 1 | | 1 |
| Museum of Vertrebrate Zoology, University of California, Berkeley | USA | 2 | | 1 |
| National Museum of Natural History, Smithsonian Institution (NMNH) | USA | 6 | | 1 |
| Royal Ontario Museum (ROM) | USA | 8 | | 1 |
| University of Michigan Museum of Zoology (UMMZ) | USA | 9 | | 1 |
| Western Foundation of Vertebrate Zoology (WFVZ) | USA | 8 | | 1 |
| Yale University Peabody Museum (YPM) | USA | 4 | | 1 |
| Kokand local museum | Uzbekistan | 1 | | |
| National Museum of Uzbekistan, ornithology collection | Uzbekistan | 8 | | |
| Samarkand State Ornithology collection (Uzbekistan) | Uzbekistan | 1 | | |

**Table S2.** Online resources regularly checked for Sociable Lapwing records between 2005 and 2020, or between 2005 and the year the platform was discontinued.

| Website | Product | Region |
|---|---|---|
| www.trektellen.nl | Online data resource | global |
| www.flickr.com | Photo repository | global |
| www.eBird.org | Online data resource | global |
| www.gbif.org | Online data resource | global |
| www.observation.org | Online data resource | global |
| www.surfbirds.com | Trip report collection | global |
| www.israbirding.com | Birdwatching website | Israel |
| www.uaebirding.com | Birdwatching website | United Arab Emirates |
| www.birdsoman.com | Birdwatching website | Oman |
| www.birds.kz | Photo repository | Kazakhstan |
| www.tarsiger.com | Birdwatching website | Western Palearctic |
| www.netfugl.dk | Birdwatching website | Western Palearctic |
| www.club300.de | Birdwatching website | Western Palearctic |
| www.orientalbirdimages.org | Photo repository | Orientalis |
| oriental bird club | Mailing list | East and South East Asia |
| African bird club | Mailing list | Africa |
| Keralabirds | Mailing list | India |
| Delhibirds | Mailing list | India |
| Indianbirds | Mailing list | India |
| Birds in Russia | Mailing list | Russia |
| RBCU list | Mailing list | Russia |
| WestPalBirds | Mailing list | Western Palearctic |
| CaucasusBirds | Mailing list | Caucasus region |
| Israbirdnet | Mailing list | Israel |

**Table S3.** Conversion of qualitative descriptions of numbers of birds into numeric estimates.

| Abundance category | Number of birds assumed |
|---|---|
| “rare” | 5 |
| “a few” | 5 |
| “singles” | 5 |
| “several” | 10 |
| “small flocks” | 10 |
| “some, some pairs” | 10 |
| “common” | 20 |
| “flocks” | 20 |
| “large flocks, large numbers” | 50 |
| “many” | 50 |
| “plentiful” | 50 |
| “few hundreds” | 500 |
| “huge flocks” | 500 |
| “gigantic flocks” | 1000 |

**Appendix S1:** Extended Acknowledgements: List of contributors to the database of sightings and participants in field surveys.

We are very grateful to the following for providing details of Sociable Lapwing sightings:

N. Abrahamsson, Sultan Al-Aseeri, Omar al-Saghier, Abdulrahman Alsirhan, Abdulmohsen Alsuraye, N.G. Aswad, John Atkins, Soner Bekir, T. Berger, Paul Bradbeer, Peter W.P. Browne, Jamie Buchan, Tom Coles, Arpit Demourari, Nihil Devasar, Jonathan Eames, Jens Eriksen, Jonathan Etzold, Omar Fadil, Wouter Faveyts, Luca Fornasari, M. Gerber, Jeff Gordon, Boris Gubin, Micha Heiß, Remco Hofland, Jon Hornbuckle, Sürreya Isfendiyaroglu, A.A. Jama, Mike Jennings, Roman Kashkarov, Mimi Kessler, Christian Ketzer, Valery Khrokov, Tatyana Kirikova, Oleg Lyakhov, Tom Martin, Phillip Meister, S. Miller, Ma Ming, Jan Peper, Mike Pope, Pedro Romero Vidal, Jonathan Rossouw, Mustafa Senocak, Ilya Smelansky, Aleksei Timoshenko, Jugal Tiwari, Rick Tovey, D. Velasco, Rick Vink, Geoff Welch, Janine Wiggers, Keith Wiggers, Simon Wotton and Vassily Zhulii.

We are very grateful to the following ornithologists and birdwatchers who participated in surveys across the range states and documented their Sociable Lapwing counts:

Ahmed Abdullah, Ross Ahmed, Ahmed Aidek, Ferdi Akarsu, Ercan Aslan, Ali Atahan, Nodir Azimov, Elizabeth Ball, Khanat Batyrkhanuly, Andrey Bazdyrev, Soner Bekir, Bahar Bilgen, Murat Biricik, Murat Bozdogan, Simon Buckell, Mazhit Buketov, Turan Çetin, Vladimir Chapurin, Mustafa Çulcuoğlu, Ahmet Demi, Mehmet Demir, Hamza Deniz, Ayhan Dursun , Mustafa Erturhan, Viktor Fedosov, Katie Field, Rob Field, Manzoor Fisher, Jeff Gordon, Olga Gordon, Andrew Gouldstone, Musa Han, Ibrahim Hashim, Geoff Hilton, Remco Hofland, Peter Iankov, Süreyya Isfendiyaroglu, Timur Iskakov, Shirin Karryeva, Roman Kashkarov, Guido Keijll, Mel Kemp, Ahman Khan, Valery Khrokov, Talgat Kisebayev, Andrew Knight, Natalya Kucheryavaya, Lars Lachmann, Oleg Lyakhov, Hywel Maggs, Mehmet Mahmutoğlu, Lyubov Malovichko, Elena Merkulova, Vladimir Morozov, Recep Mungan, Evgenii Murzakhanov, Yusuf Özbey , Korhan Özkan, Mikhail Pleshanov, Genrietta Pulikova, Alexandr Putilin, Fiona Roberts, Eldar Rustamov, Graeme Ruthven, Mohammed Salem Khan, Albert Salemgareev, Jumamurad Saparmuradov, Holger Schielzeth, Martin Scott , Ilya Smelanchev, Valentin Soldatov, Muhd Tariq, Anna Ten, Aleksei Timoshenko,

Kseniya Timoshenko, Mikail Topdas, Mukhtar Turaev, Atamurat Veyisov, Col Waseem, Graham White and John Wills.
